# The cynosure of CtBP: evolution of a bilaterian transcriptional corepressor

**DOI:** 10.1101/2022.06.23.497424

**Authors:** Ana-Maria Raicu, Dhruva Kadiyala, Madeline Niblock, Aanchal Jain, Yahui Yang, Kalynn M. Bird, Kayla Bertholf, Akshay Seenivasan, David N. Arnosti

## Abstract

Evolution of sequence-specific transcription factors clearly drives lineage-specific innovations, but less is known about how changes in the central transcriptional machinery may contribute to evolutionary transformations. In particular, transcriptional regulators are rich in intrinsically disordered regions that appear to be magnets for evolutionary innovation. The C-terminal Binding Protein (CtBP) is a transcriptional corepressor derived from an ancestral lineage of alpha hydroxyacid dehydrogenases; it is found in mammals and invertebrates, and features a core NAD-binding domain as well as an unstructured C-terminus (CTD) of unknown function. CtBP can act on promoters and enhancers to repress transcription through chromatin-linked mechanisms. Our comparative phylogenetic study shows that CtBP is a bilaterian innovation whose CTD of about 100 residues is present in almost all orthologs. CtBP CTDs contain conserved blocks of residues and retain a predicted disordered property, despite having variations in the primary sequence. Interestingly, the structure of the C-terminus has undergone radical transformation independently in certain lineages including flatworms and nematodes. Also contributing to CTD diversity is the production of myriad alternative RNA splicing products, including the production of “short” tailless forms of CtBP in Drosophila. Additional diversity stems from multiple gene duplications in vertebrates, where up to five CtBP orthologs have been observed. Vertebrate lineages show fewer major modifications in the unstructured CTD, possibly because gene regulatory constraints of the vertebrate body plan place specific constraints on this domain. Our study highlights the rich regulatory potential of this previously unstudied domain of a central transcriptional regulator.

## Introduction

The C-terminal Binding Protein (CtBP) is a transcriptional corepressor that plays critical roles in development, tumorigenesis, and cell fate (Boyd *et al*. 1993; Schaeper *et al*. 1995; reviewed in Chinnadurai 2007; Stankiewicz *et al*. 2014). CtBP has been implicated in human cancer, and is being investigated as a potential drug target (Nadauld *et al*. 2006; Barroilhet *et al*. 2013; Deng *et al*. 2013; Dcona *et al*. 2017). CtBP was first identified as a protein that binds the C-terminus of the adenoviral E1A oncoprotein and was later found to interact with diverse cellular transcription factors via their PLDLS motif, creating complexes that alter chromatin (Boyd *et al*. 1993; Schaeper *et al*. 1995; Nibu *et al*. 1998; Turner and Crossley 2001; Shi *et al*. 2003). This cofactor transcriptionally regulates genes involved in apoptosis, cell adhesion, and the epithelial-to-mesenchymal transition, functioning as a repressor in most cases, although it can directly activate promoters in some contexts (Grooteclaes *et al*. 2003; Fang *et al*. 2006; Jin *et al*. 2007; Paliwal *et al*. 2012). Unique among transcriptional coregulators, CtBP structurally resembles D-2-hydroxyacid dehydrogenases and binds the NAD(H) cofactor (Chinnadurai *et al*. 2002; Kumar *et al*. 2002). *In vitro*, CtBP proteins can use a variety of alpha hydroxyacids as substrates, but the natural *in vivo* substrate, if any, remains unknown (Kumar *et al*. 2002, Balasubramanian *et al*. 2003; Achouri *et al*. 2007). Mammalian cell culture studies have shown that the residues required for *in vitro* catalytic activity are not required for transcriptional repression or the apoptotic activities of CtBP, but in the fly, residues of the dehydrogenase active site are required for normal activity of this repressor (Grooteclaes *et al*. 2003; Zhang and Arnosti 2011).

CtBP binding to NAD(H) is necessary for its normal functions, and substantial evidence indicates that NAD(H) binding supports CtBP dimerization and tetramerization (Kumar *et al*. 2002; Balasubramanian *et al*. 2003; Sutrias-Grau & Arnosti 2004; Mani-Telang *et al*. 2007; Madison *et al*. 2013; Bellesis *et al*. 2018; Jecrois *et al*. 2021). Tetramerization has been shown to be required for transcriptional repression, as tetramer-destabilizing mutants have compromised transcriptional regulatory activity (Bhambhani *et al*. 2011; Bi *et al*. 2018; Ray *et al*. 2017; Jecrois *et al*. 2021). Additionally, it is the oligomeric form of CtBP that associates with other factors, suggesting this is the relevant form for transcriptional regulation (Shi *et al*. 2003; Jecrois *et al*. 2021). Both NAD+ and the reduced NADH cofactor promote oligomerization, but their relative binding affinities to CtBP have been disputed: one study found NADH to have >100-fold stronger binding than NAD+, while other studies indicate no differences in binding affinity (Fjeld *et al*. 2003; Zhang *et al*. 2002; Bellesis *et al*. 2018). More recently, the Boyer lab has shown through ultracentrifugation and ITC that there is only a 9-fold binding affinity difference, and that CtBP is saturated with NAD+, suggesting that it does not respond to cellular redox levels (Erlandsen *et al*. 2022). Currently, it is not clear whether NAD(H) binding to CtBP is merely a structural element, or whether the presence of the cofactor may bridge cellular metabolism and gene regulation.

CtBP proteins contain a variety of functional elements, including an N-terminal (NTD) substrate binding domain that overlaps with the conserved dehydrogenase domain, and the flexible and unstructured C-terminal domain (CTD). Differential promoter usage and alternative splicing produces distinct mammalian CtBP isoforms, with a CtBP2 RIBEYE variant having a sizable N-terminal extension that is unrelated to domains found in other CtBP isoforms (Schmitz *et al*. 2000). A hydrophobic cleft toward the N-terminus binds the E1A PLDLS motif, with additional interactions with cellular factors also mediated by the RRT-binding surface groove (Schaeper *et al*. 1995; Quinlan *et al*. 2006; Kuppuswamy *et al*. 2008). The central dehydrogenase domain includes a Arg-Glu-His (REH) catalytic triad and a Rossman fold involved in NAD(H) binding (Kumar *et al*. 2002).

The structural element that seems to distinguish CtBP proteins most clearly from more distantly related alpha hydroxyacid dehydrogenases is the unstructured CTD, which has not been structurally resolved (Nardini *et al*. 2006). This portion of the protein is the site of post-translational modifications such as phosphorylation and sumoylation (Kumar *et al*. 2002; reviewed in Chinnadurai *et al*. 2007; Jecrois *et al*. 2021). Dimeric and tetrameric forms of the protein lacking the entire C-terminal region can be obtained *in vitro*, indicating that this domain is not essential for this structural aspect of CtBP (Jecrois *et al*. 2021). One study suggested that an intact CTD was required for CtBP tetramerization, but Royer and colleagues demonstrated through SEC-MALS and cryoEM that the minimal dehydrogenase domain, without a CTD, can tetramerize in the presence of NAD(H) (Madison *et al*. 2013; Bellesis *et al*. 2018; Jecrois et al. 2021). Regarding function, the mammalian CtBP1 lacking the final 86 residues can still function as a repressor in cell culture (Kuppuswamy *et al*. 2007). Additionally, a “short” CtBP isoform is sufficient to rescue lethality of *dCtBP* loss in Drosophila, further indicating that core functions are possible in the absence of this domain (Zhang and Arnosti 2011). Thus, the roles of the CTD in oligomerization, transcriptional regulation, and other nuclear activities still remain to be defined.

Diverse CtBP proteins are found within Metazoa; invertebrates can express several isoforms from a single locus through alternative splicing and alternative promoter usage, while vertebrates have additional diversity through gene duplications that produced two or more paralogous CtBP genes. In mammals, multiple isoforms are expressed from each CtBP1 and CtBP2 paralog, which have both overlapping and unique genetic roles in the cell--both nuclear and extracellular (Katsanis and Fisher 1998; Schmitz *et al*. 2000; Hildebrand and Soriano 2002; reviewed in Chinnadurai *et al*. 2007). *CtBP2* is an essential gene in the mouse, with null mutants showing embryonic lethality. *CtBP1* null mice are viable, but exhibit developmental phenotypes (Hildebrand and Soriano 2002). In contrast, Drosophila possesses a single CtBP gene that expresses diverse CtBP isoforms through alternative splicing, affecting in particular the CTD (Nibu *et al*. 1998; Poortinga *et al*. 1998). Two major isoforms, the “long” and “short” forms, differ mainly in the C-terminus and are differentially expressed in development (Sutrias-Grau *et al*. 2004; Mani-Telang and Arnosti 2007). The long version (CtBP_L_) contains a ∼90 residue extension not found in the short protein (CtBP_S_). CtBP_S_ is the most abundant isoform in Drosophila, and it represses just as well as CtBP_L_ when tethered to Gal4 *in vivo* (Sutrias-Grau *et al*. 2004; Mani-Telang and Arnosti 2007). Loss of CtBP is lethal in Drosophila; this phenotype can be rescued by expression of either a CtBP_S_- or CtBP_L_-expressing transgene (Zhang and Arnosti 2011). However, there is an indication that expression of both isoforms is important; in this system, rescue by CtBP_L_ leads to significant changes in several target genes, not seen with rescue by CtBP_S_ (Zhang and Arnosti 2011).

The deeper biological significance of the CTD encoded in CtBP genes, and reason for its conservation, are still unknown. We hypothesize that the CTD may play a role in regulation and/or turnover, interactions with cofactors to regulate transcription, or protein localization. To lay the groundwork for experimental analysis of the CtBP CTD, we have undertaken a comprehensive comparative approach and assessed characteristics of the C-terminal portion of the protein across the animal kingdom. Investigating richly-resourced dipteran and other arthropod genomic resources and extending to invertebrates and vertebrates in general, we describe the conservation and variation found in divergent clades, pointing to likely functional aspects of this domain.

## Results

### 1) Origin of CtBP

Sequence conservation and functional similarities support the orthology of well-studied CtBP genes from mammals and Drosophila. The high degree of sequence divergence noted in the CTD of the presumed ortholog from *C. elegans* raises a question of what features most reliably support orthology in this gene family (Nicholas *et al*. 2008). A high level of sequence similarity is noted across the dehydrogenase domain (arthropod to vertebrate CtBP1 typically ∼70% identity; **Figure S1**). However, whether genes with lower sequence identities in other organisms are orthologs has not been comprehensively assessed. The Arabidopsis ANGUSTIFOLIA (AN) gene encodes a divergent homolog of CtBP that is not likely to be orthologous to animal genes; AN has a lower (∼30%; **Figure S1**) level of sequence similarity across the core dehydrogenase domain, lacks conserved catalytic residues, and has cytoplasmic functions related to microtubule regulation, membrane trafficking, and stress response (Kim *et al*. 2002; Folkers *et al*. 2002; Bhasin *et al*. 2017). This protein does not mediate repression in heterologous animal assays, although mutants show changes in gene expression (Kim *et al*. 2002; Stern *et al*. 2007; Xie *et al*. 2020). Similarly, fungal and choanoflagellate CtBP homologs have low (∼30%; **Figure S1**) levels of sequence similarity across the dehydrogenase domain, and best hits using mammalian or dipteran searches identify genes encoding proteins that are annotated as dehydrogenases, lacking any unstructured CTD.

We asked at which point in the metazoan phylogeny we could identify CtBP-encoding genes with high levels of sequence similarity, similar to those observed in initial mammalian-insect alignments. We did not identify homologs with such levels of similarity, or extended unstructured CTD, in representative genomes from Cnidaria or Porifera (**Figure S1**). In these genomes, the closest homologs exhibited levels of sequence identity similar to those of fungi and plants (∼30%). Additionally, these species lack a C-terminal domain similar to that found in flies and humans. The first CtBP gene therefore likely arose in a common ancestor to bilaterians, a decisive point in animal evolutionary history, when new combinations of gene batteries appeared that regulated novel morphological traits. As we report here, certain unique features of CtBP appear to be conserved, though not entirely, across protostomes and deuterostomes. Diversity in CtBP has been achieved over time through gene duplication in Vertebrata, and generation of alternative isoforms through transcriptional and splicing variation. To better understand the molecular processes underlying CtBP diversity, we first considered these genes in Drosophila, where extensive genomics combined with experimental work inform our understanding.

### 2) Drosophila melanogaster expresses alternatively spliced CtBP isoforms

The *Drosophila melanogaster* CtBP gene, which is essential for development, produces a number of alternatively spliced transcripts (Poortinga *et al*. 1998; Mani-Telang and Arnosti 2007). Ten isoforms have been reported, differing in their transcriptional start sites (TSS), length of their untranslated regions (UTR), and inclusion of 3’ exons. From this one genomic locus, two general types of proteins are produced: the CtBP “long” (CtBP_L_) and CtBP “short” (CtBP_S_), named based on the length of their CTDs (**Figure 1A**). CtBP_L_ isoforms are predicted to be between 473 and 481 residues in length, creating proteins of ∼50 kDa in size, and CtBP_S_ proteins are 379 to 386 residues long and produce proteins of ∼42 kDa. The long and short isoforms are generated through alternative splicing, and their existence has been corroborated by western blot, although this method does not differentiate among potential isoforms that differ solely by a few amino acids (Mani-Telang and Arnosti 2007). The ten reported mRNA isoforms are predicted to encode seven protein variants, ranging in size from 379 to 481 residues (**Figure 1B**). The CtBP_L_ isoforms incorporate the 3’ most protein coding exons, terminating in the same sequence motif; they differ in alternative splicing of three short motifs within the core and CTD (**Figure 1C**). In contrast, the ORFs of the CtBP_S_ isoforms end shortly after the catalytic core and terminate in one of two ways: reading through a splice donor site after protein coding exon 5 into the adjacent intron, or through usage of an alternative splice acceptor site 5’ of protein coding exon 7, which encodes the last portion of the CtBP_L_ isoforms in a different reading frame. In addition, these short isoforms also differ in the retention or deletion of a short VFQ tripeptide found near the start of the CTD (**Figure 1D**). The remaining NTD and core sequences are identical among CtBP isoforms in the fly. Of interest is the clear difference between the short and long isoforms, which we have shown to differ in their spatial and temporal expression in *D. melanogaster*, and which may play unique roles in development (Zhang and Arnosti 2011). In fact, the two major isoforms are developmentally regulated, and CtBP_S_ is believed to be the predominant form expressed across development (Mani-Telang and Arnosti 2007; Zhang and Arnosti 2011).

**Figure 1.**
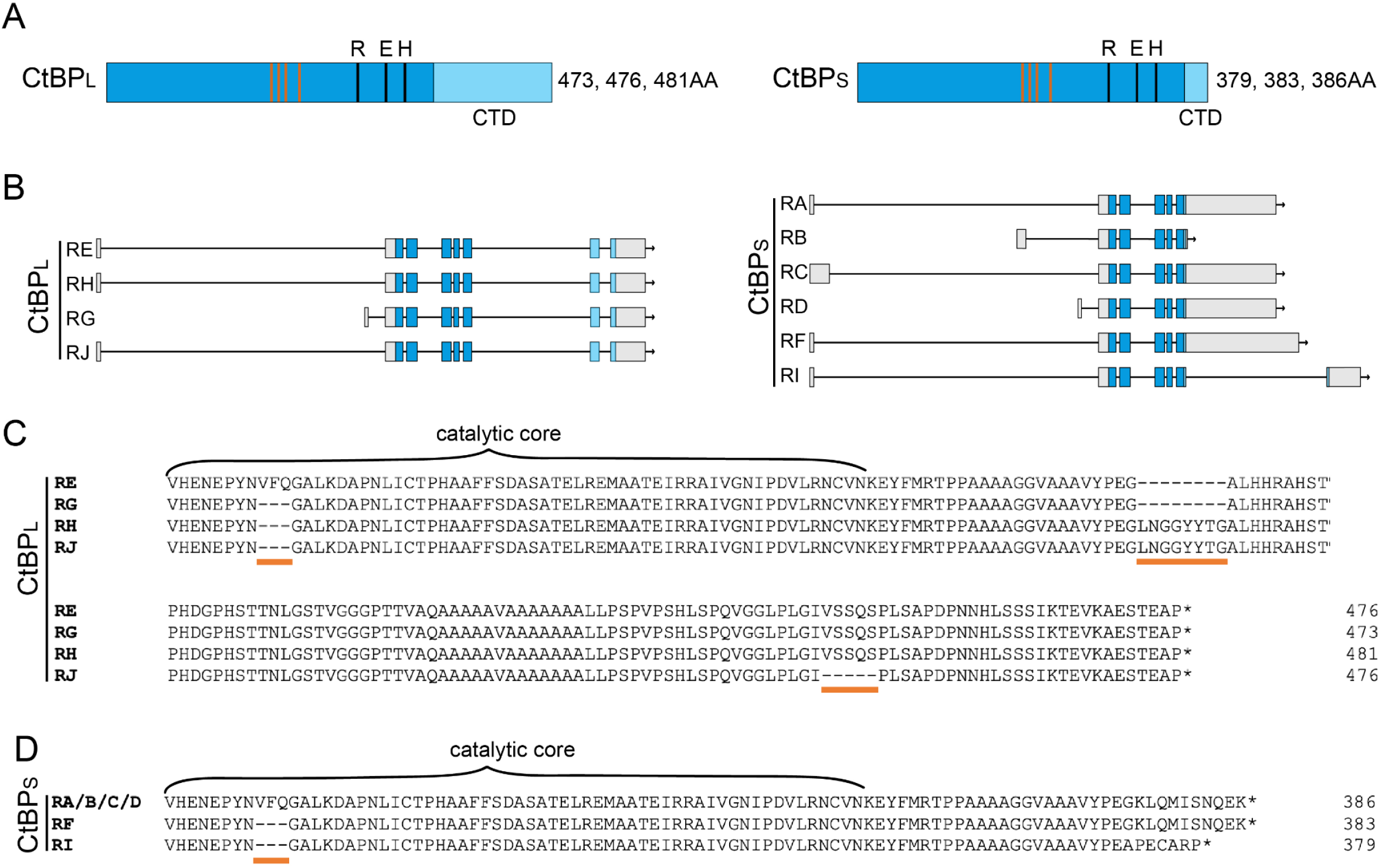
The CtBP locus in *Drosophila melanogaster* produces variant transcripts and proteins with different C-terminal lengths. **A)** Two general types of CtBP proteins are produced in *D. melanogaster:* long (CtBP_L_) and short (CtBP_S_). The proteins are almost identical in their N-terminus and central catalytic domain (blue), and differ in the sequences and lengths of the C-terminal domain (light blue). Proteins of three different sizes are predicted to be produced for each isoform. Orange vertical bars indicate the four residues involved in NAD binding, and black lines indicate the three residues making up the catalytic triad (REH). **B)** Schematic representation of the 10 transcripts produced from the CtBP locus. Gray boxes indicate 5’ and 3’ UTRs, blue boxes indicate protein-coding exons, and horizontal lines are introns. Isoforms E, H, G, J encode long versions of the protein and use two different TSSs. Isoforms A, B, C, D, F, and I encode short versions of the protein and use three different TSSs. **C)** Alignment of the C-terminal region of the conserved core and the CTD indicates that four different long proteins are encoded, which differ with the inclusion or deletion of three small motifs in the core and CTD: a VFQ tripeptide, an LNGGYYTG motif, and VSSQS motif (orange horizontal bars), all of which are spliced out of the mRNA in different combinations. **D)** Alignment of the CTD of the short isoforms indicates that three different proteins are predicted to be produced, which differ with the inclusion or deletion of the same VFQ tripeptide and terminate with SNQEK or APECARP.

### 3) Long and short CtBP isoforms are conserved throughout Drosophila

We next assessed the potential conservation of the variant CtBP isoforms produced through alternative splicing in additional Drosophila species, which may point to functional roles of these products. We examined predicted protein sequences of CtBP isoforms from eleven additional drosophilids (**Figure 2A**). For every species examined, multiple mRNA sequences indicate that both long and short isoforms are encoded, and that they differ in the retention or loss of the same short segments encoding VFQ, LNGGYYTG, and VSSQS that were observed in *D. melanogaster* (data not shown). The conservation of these variants suggests that expression of long and short isoforms of CtBP, as well as the exclusion/retention of short motifs, are functionally important. Considering the CtBP isoforms encoded in the twelve drosophilid mRNAs, the NTD and catalytic core sequences are highly conserved. In contrast, CTD sequences themselves show more evolutionary variation, particularly in the center of the CTD, with presence or absence of alanine- and proline-rich sequences (**Figure 2B**). All species express mRNAs encoding CtBP short proteins ending in AP/SECARP, using a conserved alternative splice acceptor site. Some also are found to produce mRNAs that create short isoforms terminating in SNQEK by reading through a splice donor site into the next intron. Additional short endings, created through alternative splicing, are seen in some species (**Figure 2C, S2A**). Surprisingly, the nucleotide sequence encoding the terminal SNQEK derived from the 3’ end of exon 5 and adjacent intron is 100% conserved in all species, which is a much higher level of conservation than noted for other coding regions, which harbor mostly synonymous changes (**Figure S2B**). We hypothesized that the absolute conservation may reflect an RNA structure that would influence the use of this splice donor site to produce a short or long isoform. We predicted the structure of this conserved sequence of RNA using the RNAstructure software (Xu and Matthews 2016), and found that the splice donor site that is used to create CtBP_L_, but is suppressed for CtBP_S_, folds into a hairpin that may sequester the GU donor site in a stem loop structure (**Figure S2C**). Although mRNA isoforms encoding CtBP_S_ terminating in SNQEK are not reported in all of these species, the absolute conservation of this specific portion of the intron suggests that the capacity to generate these isoforms is conserved. Other Drosophila genes have been shown to exhibit a high level of conservation in certain intronic regions that can form RNA hairpin structures to influence alternative splicing events (Raker *et al*. 2009). In contrast, the nucleotide sequence encoding the AP/SECARP short form variants is not as highly conserved, with both synonymous and nonsynonymous substitutions present in the protein coding exon, and high divergence in the preceding intron (data not shown). This indicates that RNA secondary structure is not important for this canonically spliced isoform. In summary, although the CTD region of CtBP is more evolutionarily variable than the core dehydrogenase domain, it is likely that the diversity of CTD structure is an important aspect of CtBP proteins in Drosophila.

**Figure 2.**
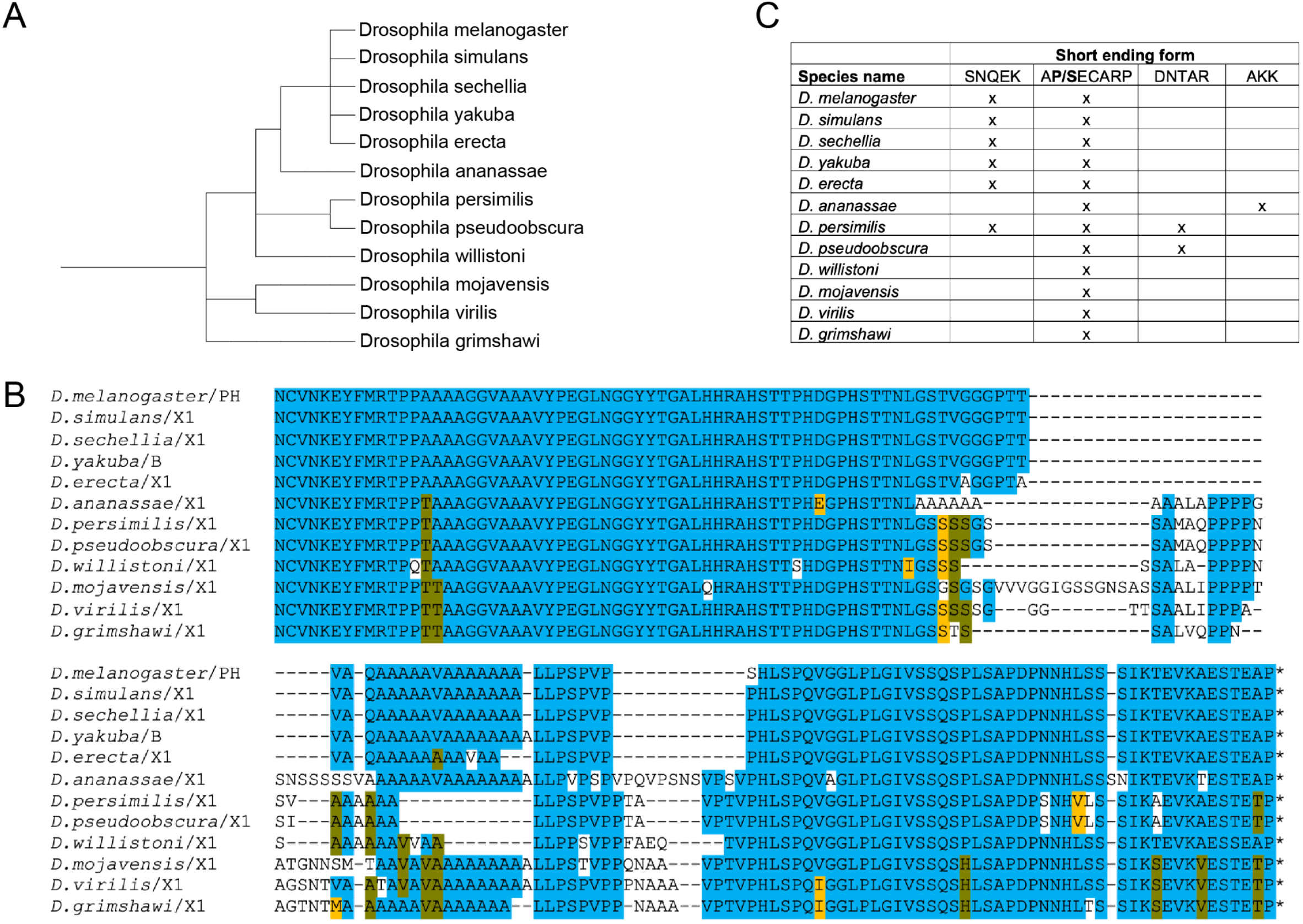
Long and short CtBP isoforms are expressed in Drosophila. **A)** Phylogenetic relationship of the Drosophila species used. These species diverged from their last common ancestor ∼40 mya. **B)** Alignment of the CTDs of the longest Drosophila CtBP_L_ sequences. Alignment begins with NCVN, which is the end of the conserved core. In this and subsequent figures, blue highlighting is used for conservation of a residue in >50% of species, gold for chemically conserved residues, and army green for conservation of a second residue in 25-50% of species. Orange horizontal bars highlight variable regions rich in proline and alanine. Specific isoform letters or numbers are indicated after the species name, and the final asterisk indicates the STOP codon. **C)** Presence (X) of various alternative CtBP_S_ terminal sequences. The SNQEK version is created by suppressing a splice donor site and extending the ORF into the intron. APECARP, DNTAR, and AKK are created through exon skipping and alternative splicing. All species express an APECARP short version and all species have the ability to encode the SNQEK version, but it is not always detected in cDNA sequences.

### 4) Conservation of long and short forms of CtBP in Diptera

Drosophila are members of the suborder Brachycera, also commonly referred to as “higher Diptera.’’ Nematocera, or “lower Diptera,” include mosquitoes, for which there are also extensive genomic resources. To assess CTD structure across Diptera, we selected thirteen higher Diptera and nine lower Diptera to perform sequence alignments. All Diptera express CtBP_L_ isoforms and many also express short variants (**Figure 3A**). Two regions exhibit a higher level of conservation among the CTD long forms; a “Central Block” containing a motif featuring hydrophobic and aromatic residues (YSEGINGGYY) with an adjacent H/S/T-rich sequence (AHSTTPHD), and a “Terminal Block” rich in prolines, followed by a short stretch of N/H and acidic residues in a conserved PExSEVH/Q terminus. It is apparent that the Drosophila C-terminal sequence noted above (ESTEAP) is a derived feature within this genus (found also in the closely related *S. lebanonensis* sequence; **Figure S3A**), although it shares the acidic character with the consensus SEVH ending found across Diptera. Among this set of sequences from higher Diptera, Drosophila also stands out for the central block of poly-alanine repeats not present in other species (**Figure 3B**). Within specific species, variants exist in which the YSEGINGGYY, AHSTTPHD, and VSSKS motifs are alternatively spliced out, as was observed in Drosophila (data not shown).

**Figure 3.**
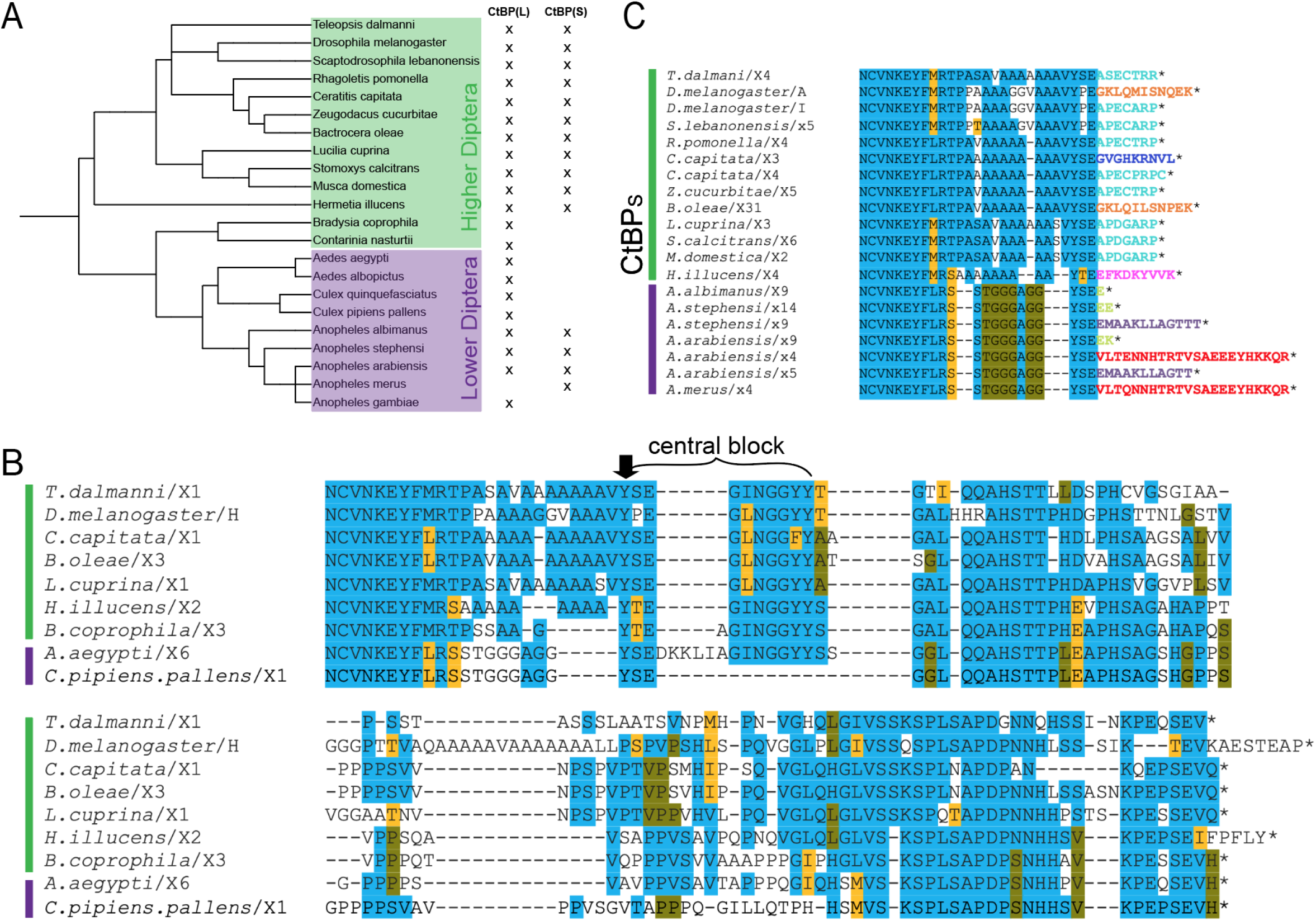
Dipterans express both long and short isoforms, with a diversity of short forms in higher and lower Diptera. **A)** Phylogenetic tree of higher and lower Diptera. Presence (X) of long or short isoforms.**B**) Alignment of CtBP_L_ in representative higher and lower Diptera. Sequences from only two mosquito genomes are presented in this alignment, as isoforms from most other species encode novel sequences at the very C terminus (**Figure S3D**). The “central block” is indicated, as well as the conserved aromatic Y (black arrow). **C)** Alignment of CtBP_S_ isoforms in Diptera. Higher Diptera encode the conserved APECARP and SNQEK, while lower Diptera express short forms not seen in their close relatives. Colors for terminal residues indicate the diversity of short endings observed in Diptera (i.e the variant ending in APECARP is in 9 of the species, colored in light blue). Variants that are found in more than one species are colored the same.

Within lower Diptera, the terminal sequences of the CtBP_L_ isoforms are highly variable--much more so than seen within higher Diptera. Across the three mosquito genera we sampled (Aedes, Culex, and Anopheles), we note many genus-specific sequences, as well as some that match those found in higher Diptera (**Figure 3B, S3D**). Depending on the species, the CtBP gene can give rise to up to six potential long CtBP isoforms and five potential short CtBP isoforms, all created through alternative splicing (**Figure S3B**). The function of each of these CtBP isoforms in mosquitos has not been experimentally assessed, but we hypothesize that the diverse proteins may serve tissue- or temporal-specific functions.

In most of the higher Diptera, we find that the CtBP_S_ isoforms SNQEK and APECARP (conserved in drosophilids) are also expressed, with the predominant short form ending in APECARP (**Figure 3C**). Interestingly, we find that splice donor site suppression occurs in *B. oleae* to form an SNQEK-like ending, as seen with *D. melanogaster*. The APECARP endings in the other Diptera are also created through alternative splicing. In contrast, only four of the sampled mosquitoes report short CtBP isoforms, all within the Anopheles genus (**Figure 3A, S3B**). These do not resemble the conserved higher dipteran SNQEK or APECARP variants, but instead have one or more of seven different short variants (**Figure S3B, C**). In summary, the production of short-and long-CtBP isoforms is found in both higher and lower Diptera, with certain sequences of the long forms showing strong conservation.

### 5) Deep conservation of arthropod CtBP structure, with lineage-specific modifications

We compared CTD sequences from CtBP genes across representative insect orders as well as from springtails, a related hexapod (**Figure 4A**). Hexapod CTD sequences exhibit a deeply conserved central block including the YPEGINGGYY and AHSTTPHD motifs, as well as a proline-rich terminal region ending in SEVH (**Figure 4B**). The ancestral SEVH-like terminal region is conserved across all sampled insect orders other than Hymenoptera, which instead feature a glycine- and proline-rich terminal sequence unique to this order. Interestingly, the springtail (*F. candida*) CTD terminates just beyond the conserved central block, highlighting two lineages in the hexapods for which the terminal regions have been remodeled. Lineage-specific “spacers” rich in alanine, glycine, and proline separate the conserved central block from more N- and C-terminal residues in Blattodea, Hemiptera, Lepidoptera, Diptera, and Hymenoptera (**Figure 4B**). CtBP_S_ isoforms are not unique to Diptera; hymenopteran isoforms also encode putative short variants (**Figure 4C**). Within Hymenoptera, alternative splicing produces the conserved order-specific RLSSRC short terminal sequence. The production of short variants appears to have arisen independently in these two orders.

**Figure 4.**
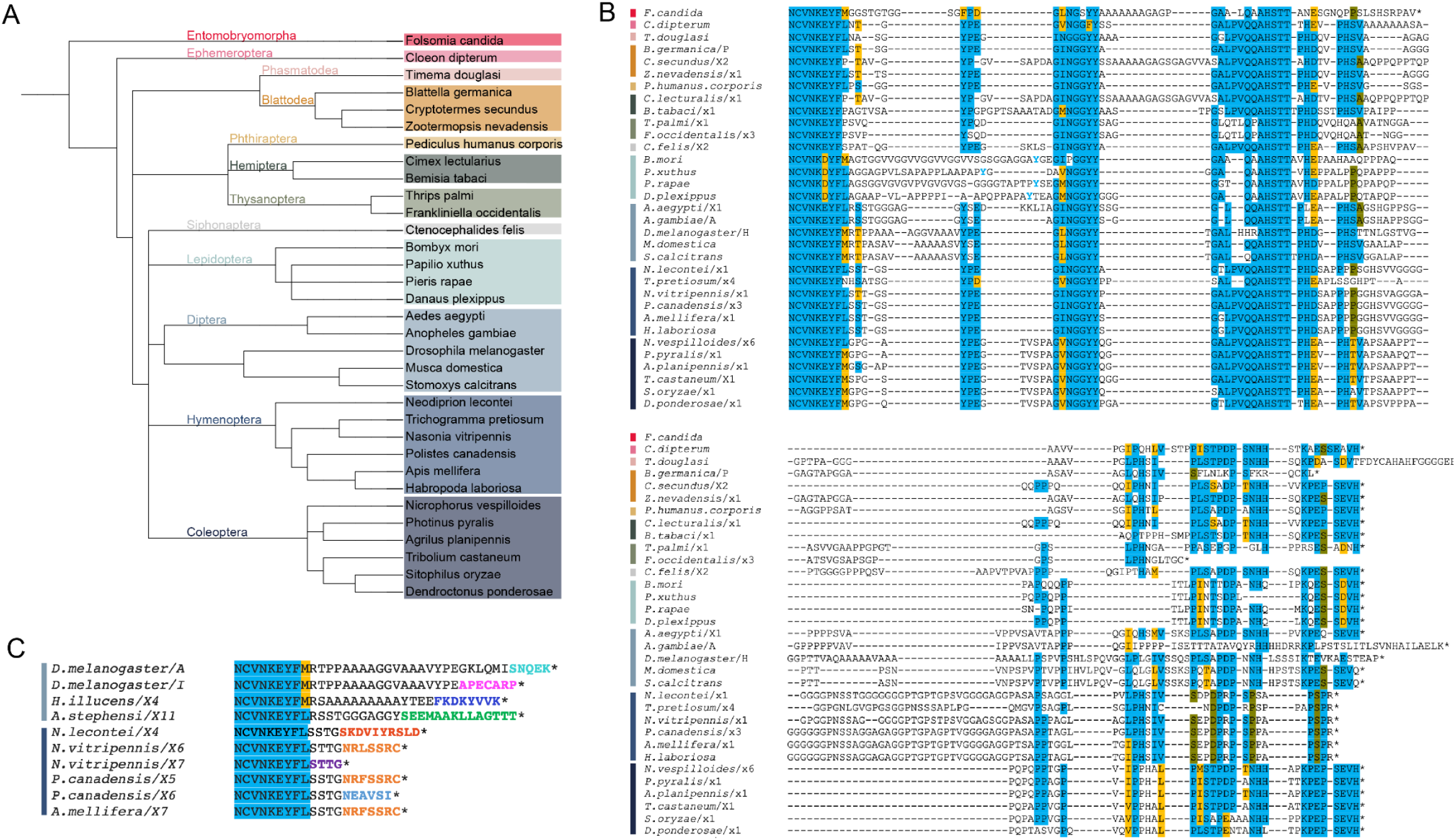
Hexapods express long forms of CtBP, with certain lineages producing short variants. **A)** Phylogenetic tree of all species analyzed in Hexapoda. Colored vertical bars in B and C correspond to orders indicated in A. **B)** Alignment of long CTDs from all Hexapod species analyzed. Certain motifs are conserved across all orders. In Hymenoptera, YG/S/TE residues within a variable region are highlighted in blue lettering to indicate presumed conservation. The *T. douglasi* sequence extends another 47 residues past what is shown. **C)** Alignment of short CTDs from Diptera and Hymenoptera indicates that within Hymenoptera, some short endings are conserved, but are distinct from sequences of Diptera. Other Insecta orders do not have short endings. Variants that are found in more than one species are colored the same (i.e the variant ending in SSRC is found in both *N. vitripennis* and *A. mellifera*, colored in orange).

A comparison of conserved hexapod sequences with those of crustaceans, myriapods, and chelicerates, which altogether make up the arthropod phylum, reveals that the central and terminal conserved regions noted in hexapods are generally conserved across arthropods (**Figure 5**). Sequences from representative species from these four groups demonstrate that four key motifs (NCVNKEY followed by an aromatic, ΨNGGYY (central block), AHSTT, and PEPSEVH) are present in all lineages, indicating that they are derived from an ancestral CtBP. From the two myriapod genomes available, no large deviations from the consensus are found. However, in crustaceans (shrimp, barnacle, and planktonic crustaceans), considerable variation is found in terminal sequence regions for all three classes analyzed (**Figure 5B, S4B**). The ancestral SEVH terminus is found only in *P. pollicipes* (Gooseneck barnacle).

**Figure 5.**
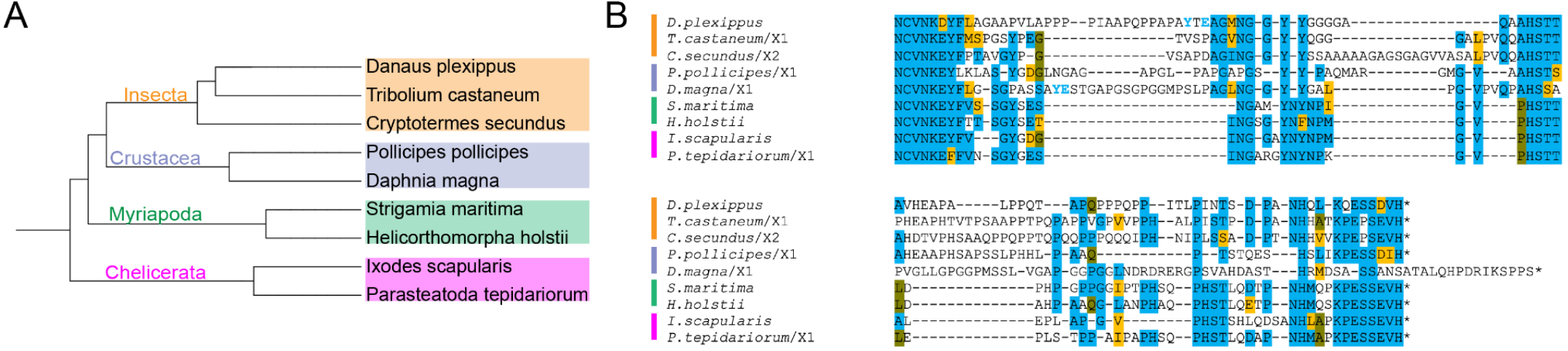
Arthropod CTDs contain conserved motifs including ancestral ending. **A)** Phylogenetic tree of representatives of the four major arthropod groups. **B)** Alignment of representative species illustrates that motifs seen across Diptera are conserved within these groups of arthropods. Vertical bars on the left represent the lineages in A. The tyrosines in light blue (found in *D. plexippus* and *D. magna*) indicate there is a conserved aromatic residue found between the NCVN and NGGYY motifs, but spaced slightly differently in these two species.

An interesting finding comes from consideration of chelicerate CtBP sequences. Most CtBP CTD sequences from this subphylum, which includes mites, ticks, scorpions, spiders, and horseshoe crabs, have clearly alignable motifs in central and terminal regions (**Figure S4D, 5B**). All the species sampled (**Figure S4C**) have only one long version of the CTD, with no indication that short isoforms are produced. For most species, very few differences in the CTD sequences are present; the conserved blocks are not separated by repeat expansions noted in some insect orders, and sequences terminate with the same SEVH motif observed in the hexapods (**Figure 5B**). Three species from the order Mesostigmata, which includes predatory and parasitic mites, share a CTD that is entirely dissimilar to other arthropod sequences (**Figure S4E**). The proline/alanine-rich CTDs of *V. destructor, V. jacobsoni*, and *G. occidentalis* do not show compelling similarity to other chelicerate sequences, including those of more distantly related ticks (Ixodida) and dust mites (Sarcoptiformes). Thus, it is evident that the CTD of CtBP has undergone a wholesale replacement in the Mesostigmata lineage, which diverged from the order Ixodida ∼300 million years ago (Mans *et al*. 2016). The novel CTD is likely to be similarly disordered, based on sequence composition, but functional properties may have changed.

### 6) Diversification in Protostomia

Within other ecdysozoan lineages, the CtBP CTD of the velvet worm (Onychophora) was substantially similar to the consensus ancestral arthropod sequence (**Figure 6A, B**). Similarly, the CTD from the priapulid *P. caudatus* provides another example of a non-arthropod ecdysozoan with highly similar CTD (**Figure 6A, B**). However, wholesale changes were found for the tardigrade CTD, where clear homology ends just after the start of the conserved central block. This alternative CTD features poly-asparagine and multiple poly-alanine stretches to generate a sequence slightly longer than those in many arthropods (**Figure 6A, B**). In Nematoda (roundworm), multiple, lineage-specific forms of the CTD were identified that bore no close similarity to the previously identified conserved elements in arthropods (**Figure S5, 6A**). Notably, the NAD-binding core of these proteins showed high conservation with invertebrate sequences (∼60%), indicating that evolutionary changes are focused on the CTD. Sequence alignments from ten roundworm species from the orders Rhabditida and Trichinnelida showed at least three distinct primary structures (**Figure S5**). In the nematodes, aside from the Caenorhabditis worms, an aromatic residue (F) universally found near the N-terminal portion of the CTD is also present. While the Caenorhabditis worms lack this feature, they have an LNMGF motif that is present at approximately the same position as the conserved central block in arthropods. In short sequence blocks, some level of similarity is present in Rhabditida, especially within Caenorhabditis (**Figure S5B**). Only CtBP_L_ isoforms were identified in the nematodes, with the Trichinella species having the longest CTDs (when compared to all other Ecdysozoa), ranging from 200 to 720 amino acids, in some cases virtually doubling the size of the CtBP protein.

**Figure 6.**
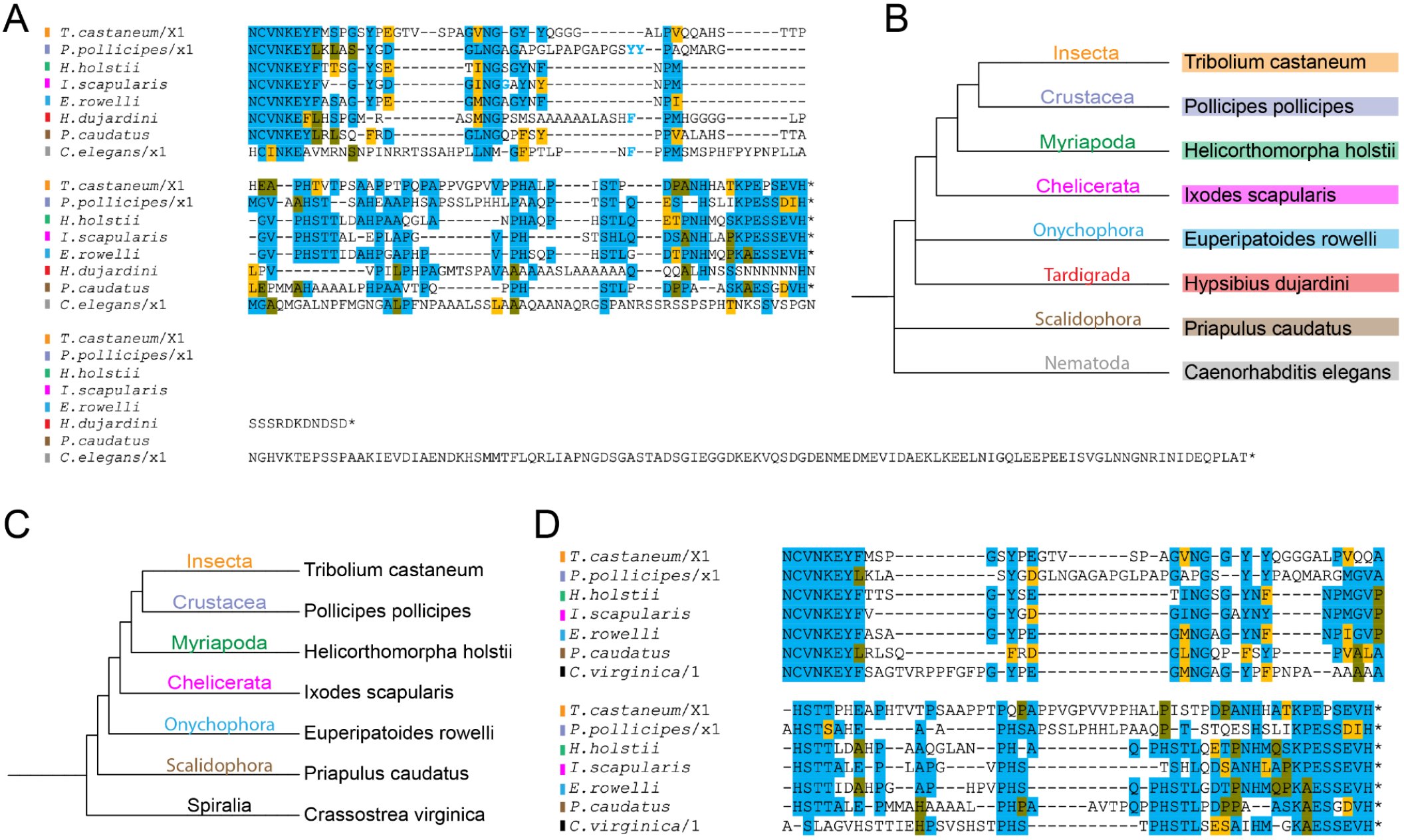
Comparative CtBP CTD alignments across Ecdysozoa and Protostomia. **A)** Alignment of CtBP sequences from representative Ecdysozoa shows conservation of central block and C-terminal sequences in most lineages. Unique and completely divergent sequences are found in tardigrades and nematodes. The residues in light blue indicate there is a conserved NGGYY-like sequence, but spaced slightly differently in these species. **B)** Phylogenetic tree of representative ecdysozoan species used in panel A. **C)** Phylogenetic tree of representative species from Protostomia (Ecdysozoa and Spiralia) that have a canonical CTD with conserved motifs. **D)** Alignment of representative protostomes shown in C illustrates the conservation of particular motifs across these invertebrates, including the central block and most C-terminal portion of the CTD.

To better understand what structural features of the CtBP CTD may be generally conserved in protostomes, we examined CtBP sequences from the morphologically diverse clade Spiralia, including molluscs, annelids, flatworms, and other taxa (**Figure S6A**). Although the adult body plans are very diverse, they share some early embryonic developmental programs, such as canonical spiral cleavage in the embryo. We selected representative species from six phyla, and found that species from all, excepting Platyhelminthes, share core conserved motifs found in the consensus ecdysozoan CTD (**Figure S6D**). A striking exception was found in certain annelids; the leeches (Hirudinea) lack a C-terminal extension entirely (**Figure S6E**). This represents the only animal lineage that appears to lack a long-form of CtBP. Polychaete annelids express CtBP with a CTD containing homology to the central block and terminal core conserved motifs, while earthworms (*L. rubellus* and *E. fetida*) and the oligochaete *O. algarvensis* bear shorter CTD sequences with small regions of sequence similarity to the protostome consensus (**Figure S6E**). Representatives of Nemertea (*N. geniculatus*; ribbon worm) and Phoronida (*P. australis;* horseshoe worm) showed strong conservation in central and terminal sequences, with minor variations (**Figure S6D**). Interestingly, the rotifer *A. vaga* CTD has recognizable homologies through the central block, and then is sharply divergent from other protostomes, a pattern of variation resembling that of the order Hymenoptera in insects (**Figure S6D, 4B**). The Platyhelminthes have unique proline and poly-alanine rich CTD sequences that do not resemble those of other species. Interestingly, there is considerable diversity within the Platyhelminthes phylum; there are weakly alignable blocks within trematode, cestode, and monogeneid CTDs while CtBP CTD sequences of triclad planaria form a separate homology set (**Figure S6B, C**). Platyhelminth CTDs range in size from 150 to 550 residues, formed by addition of novel residues to the terminus.

Overall, deep conservation of the CTD of CtBP within Protostomia is punctuated by replacement of the ancestral sequence with an alternative in multiple lineages (**Figure 6A, D**). The chemical nature of the typical CTD sequence, as well as the sizes, are generally conserved (**Figure S10A, B**). Only the leech appears to have done away with the CTD entirely, but various arthropods have devised splice variants that presumably allow for facultative expression of a short form, as in Drosophila. The significance of massive alternative CTDs (>500 residues), as seen in nematodes and flatworms, remains obscure.

### 7) Conservation of CTD sequences between protostomes and deuterostomes

In contrast to a single CtBP gene found across Protostomia, mammals have two CtBP paralogs, CtBP1 and CtBP2, which have both overlapping and unique functions in development (Katsanis and Fisher 1998; Hildebrandt and Soriano 2002). *CtBP1* null mice are viable but have developmental phenotypes, while *CtBP2* null mice are embryonic lethal (Hildebrandt and Soriano 2002). The human CtBP1 and CtBP2 proteins exhibit ∼90% conservation in the dehydrogenase core, with most of the remaining 10% reflecting chemically conserved substitutions. Interestingly, the CTD itself has more variation, with only 50% of the primary sequence being conserved between the two paralogs (**Figure S7A**). We can conclude, however, that the CTD sequences are derived from a common ancestor. Specific motifs in the CTD show stronger conservation, such as a central PELNGAxYRY motif, as well as the aromatic W situated between the N-terminal region and this motif, both of which resemble conserved motifs of the invertebrates (**Figure S7A, 6D**). The charged residues (one basic/two acidic) at the very terminus also appear to represent conserved features, while various alignable prolines are less compelling as evidence of homology for these overall proline-rich sequences. Deeply conserved AHSTT and PHS-PHS motifs located between the central motif and terminus in many protostomes are not conserved in the human CTDs (**Figure S7B**), but the overall length of the CTDs (∼100 residues) is similar to those of representative protostomes (**Figure S10A**). Overall, the similarities argue for a common CTD sequence shared by the last common ancestor of mammals and invertebrates, a feature that was not apparent when only a few CtBP genes were known, including the *C. elegans* CtBP with its derived CTD.

To better understand evolutionary processes in deuterostomes, we turned to genomes of species representing echinoderms, acorn worms (hemichordates), and non-vertebrate chordates, including tunicates (urochordates) and lancelets (cephalochordates) (**Figure S7D**). In contrast to mammals, only a single CtBP gene is found in these species, as in protostomes. These CtBP CTDs are clearly homologous; they share a lone tryptophan toward the N-terminus, adjacent to the central block PELNGxYRY (similar to central block from protostomes), more lineage-specific blocks, and a highly conserved terminus with one basic and two acid residues in conserved spacing (**Figure S7C**). Insertions between these more conserved regions are present in the two tunicate CtBP sequences, generating a longer CTD.

### 8) Vertebrates encode multiple CtBP genes

To better understand the molecular transformations that occurred as vertebrates diversified from the last common ancestor of these distant relatives, we analyzed CtBP isoforms in Vertebrata, where multiple rounds of whole genome duplications have increased the number of many paralogous genes (**Figure 7A**). In vertebrate genome annotations, paralogous genes are based on presumed similarities to mammalian CtBP1 or CtBP2 paralogs; however, we find that these designations are in some cases inaccurate, based on the presence of highly conserved residues characteristic of one or the other paralog. We prepared a systematic set of criteria to reliably designate a paralog CtBP1-like, CtBP1a, or CtBP2-like (see Materials and Methods; **Figure 7B**).

**Figure 7.**
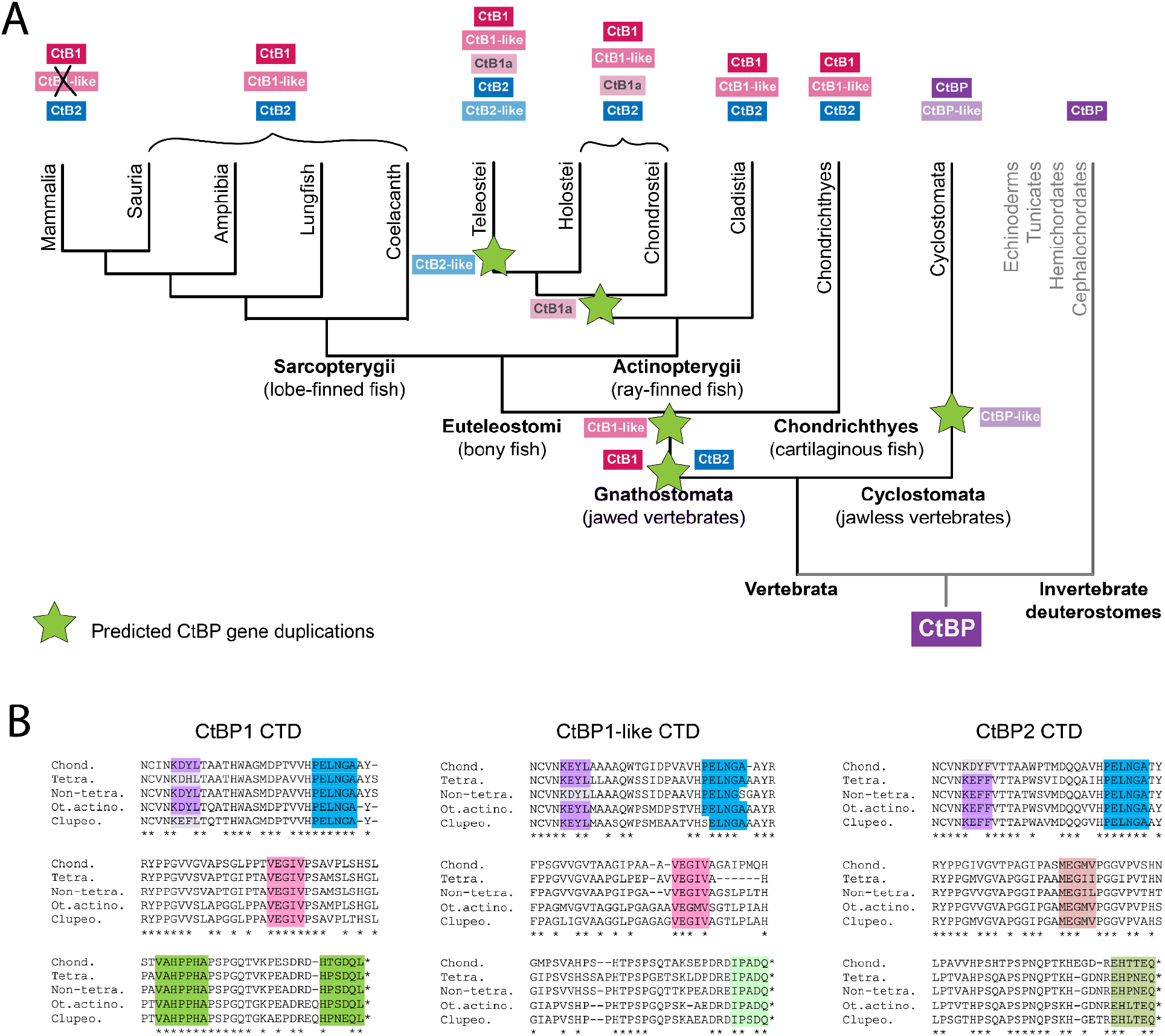
CtBP paralogs in vertebrates. **A)** All vertebrates express two or more CtBP genes. Based on sequence similarities, an independent duplication (green star) is suggested to have happened in Cyclostomata, whereas three paralogs (CtBP1, CtBP1-like, and CtBP2) originated in an ancestor to jawed vertebrates, possibly generated through whole genome or independent gene duplication events. Loss of the CtBP1-like gene occurred only in mammals, while additional gene duplications occurred at different times in ray-finned fish. **B)** Alignment of representative Gnathostomata sequences of the CtBP1, CtBP1-like, and CtBP2 CTDs. The sequences do not represent those of a particular species, but rather a consensus that illustrates the representative CTD for that particular clade. For all alignments: Chond. (Chondrichthyes); Tetra. (Tetrapods); Non-tetra. (Non-tetrapod sarcopterygians, including lungfish and coelacanth); Ot. actino (Other actinopterygii, includes Cladistia, Chondrostei, Holostei and non-Clupeocephala Teleostei); Cupleo. (Clupeocephala, includes some Teleostei like zebrafish, pufferfish, and northern pike). Asterisks on the bottom indicate complete conservation of a particular residue. Purple highlighting indicates a region characteristic of a particular paralog grouping, and light purple indicates a lineage-specific derivation. Blue highlight indicates a sequence that’s conserved across all clades, across all paralogs. Pink highlight indicates a motif unique to CtBP1 and 1-like, but that differs in CtBP2. Green highlight indicates a motif that is highly conserved within each protein family, and is representative of that protein, but not of the other paralogs.

In the lamprey (Cyclostomata, a jawless fish), two paralogs are found which we have named CtBP and CtBP-like (**Figure 8**). The lamprey CtBP is more similar to the vertebrate CtBP1 and CtBP2 than to its own paralog, showing ≥85% identity to the vertebrate proteins across the dehydrogenase core. The lamprey CtBP and vertebrate CtBP1 and CtBP2 CTD sequences are likewise very similar. In contrast, the lamprey CtBP-like CTD, while clearly derived from the canonical ancestral sequence, is less similar, and contains insertions between conserved blocks. This evidence suggests that CtBP-like is derived from an independent duplication of the single CtBP gene in jawless fish.

**Figure 8.**
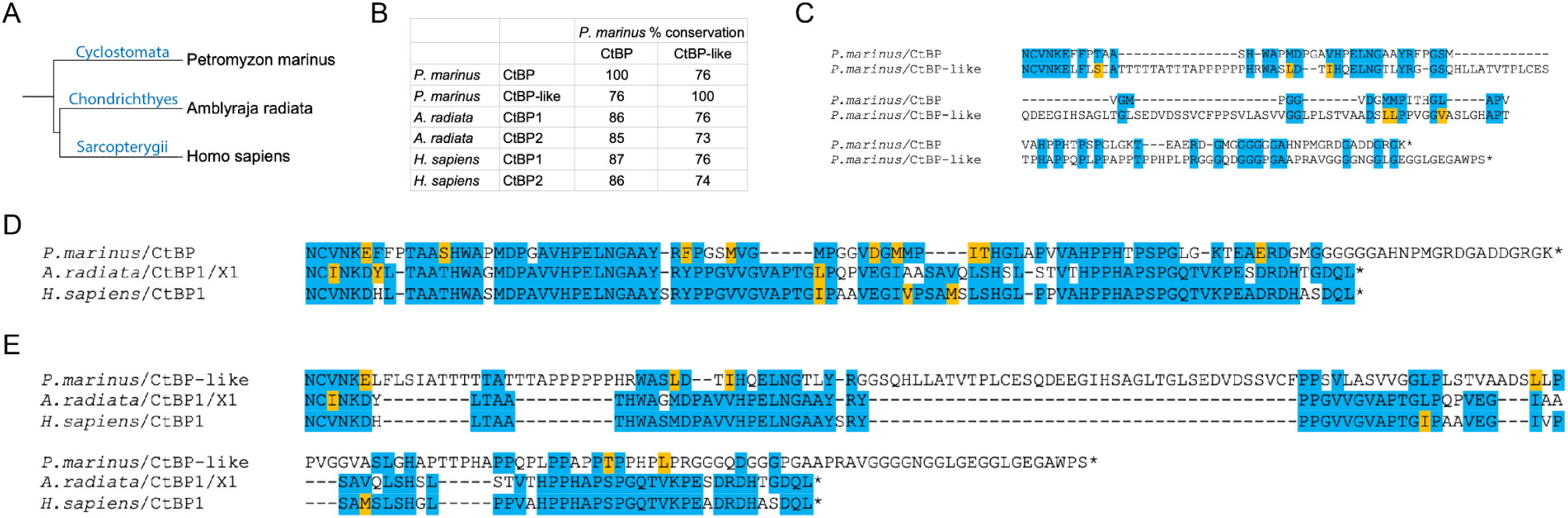
Cyclostomata CtBP CTD sequences differ from those of other deuterostomes. **A)** Phylogenetic tree representing the relationship between the lamprey (Cyclostomata, basal vertebrates), thorny skate (Chondrichthyes), and human (Sarcopterygii). **B)** Comparison of the percent conservation of the dehydrogenase core of the lamprey (*P. marinus*) CtBP and CtBP-like to CtBP1 and CtBP2 from a representative Chondrichthyes (*A. radiata)* and Sarcopterygii (*H. sapiens)*. Numbers indicate percentage of completely conserved residues including and between the RPLVALL and NCVN motifs. The lamprey CtBP has a higher degree of similarity to vertebrate CtBP1 and CtBP2 than does the lamprey CtBP-like paralog. CtBP-like may have originated as a duplication specific to the lamprey, and then diverged within this lineage.C) Alignment of the lamprey CtBP and CtBP-like CTDs indicates low conservation, and differences in CTD length. **D)** Alignment of the lamprey CtBP CTD with that of representative vertebrates’ CtBP1 CTD. The terminal residues of lamprey CtBP show a derived extension, with otherwise high level of similarity. **E)** Alignment of the lamprey CtBP-like CTD with representative vertebrates’ CtBP1 CTD. Residues more C-terminal to the central block motif constitute a much longer sequence that appears to be derived in this lineage.

In contrast, species of cartilaginous fish (Chondrichthyes) have CtBP1, CtBP1-like, and CtBP2. The CtBP paralogs in this ancient fish lineage may have originated during basal whole genome duplication events in the jawed fish (Gnathostomata) (**Figure 7A, 9**). The CTDs from Chondrichthyes are similar to the lamprey CtBP CTD, but differ greatly from the lamprey CtBP-like CTD (**Figure 8D, E**). Chondrichthyes are the first lineage in which we find expression of three different CtBP proteins, with conservation of the third, CtBP1-like, across the selected species. Interestingly, short isoforms of CtBP1 and CtBP2 are reported in some of the species, suggesting that formation of a CtBP_S_ isoform arose independently in these vertebrates, similar to what was observed in certain insect orders (**Figure 9B**).

**Figure 9.**
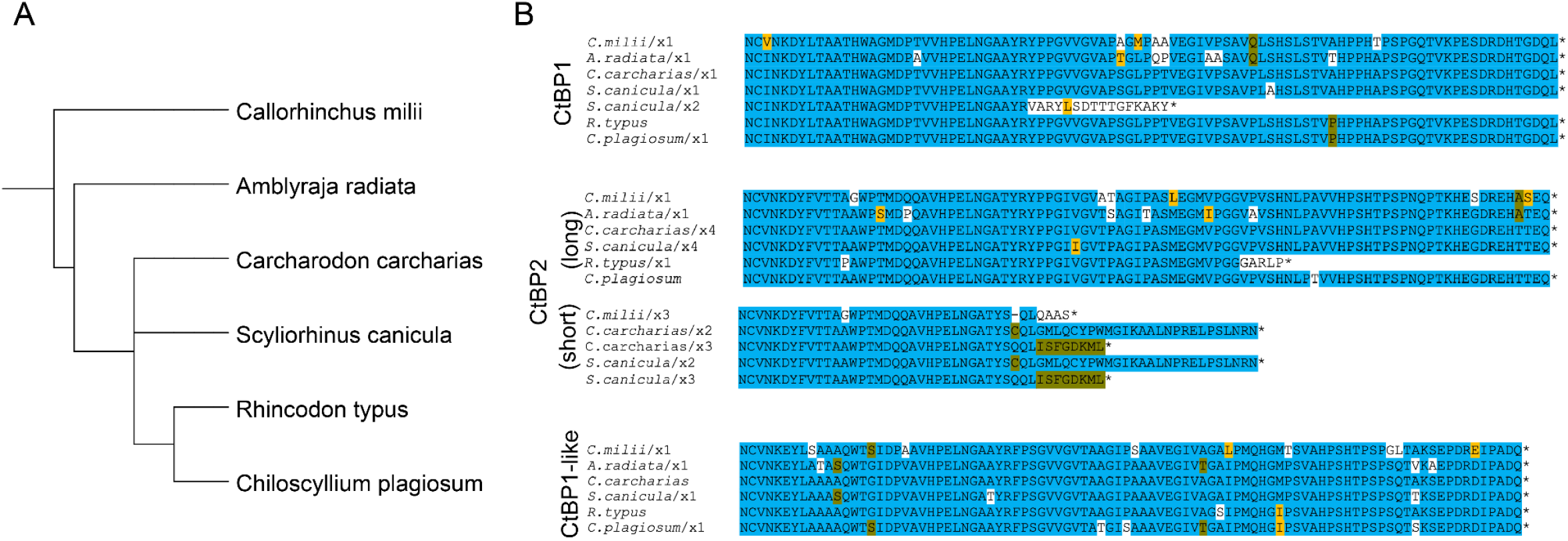
Chondrichthyes encode three CtBP genes. **A)** Phylogenetic tree of selected Chondrichthyes. **B)** Alignment of representative isoforms from each of six cartilaginous fish indicates that the CTDs are very highly conserved within CtBP1, CtBP2, and CtBP1-like, with short isoforms appearing in both CtBP1 and CtBP2.

An examination of representative species from the two groups of bony fish (Euteleostomi) reveals additional changes in CtBP gene copy number. In the lobe-finned fish (Sarcopterygii), extant species have homologs of CtBP1, CtBP1-like, and CtBP2 as found in the ancestral Chondrichthyes, with retention in most tetrapods. The CtBP1-like paralog is lost solely in mammals (**Figure 7A**). In ray-finned fish (Actinopterygii), additional CtBP1a and CtBP2-like genes are found. Teleost-specific gene duplication events are associated with up to five CtBP genes in certain lineages (**Figure 7**).

Characteristic residues present in CtBP1, CtBP1-like, and CtBP2 across Gnathostomata reveal particular segments of the CTD that have undergone modifications at different evolutionary times. For instance, more recent derivations are represented by a tetrapod-specific change in the more N-terminal portion of the CtBP1 CTD from the ancestral KDYL to KDHL, while the same region underwent a conversion from KDYL to KEFL within Clupeocephala, a specific clade within Actinopterygii (purple highlight; **Figure 7B**). A more ancient derivation is observed in a comparable location in the CtBP2 CTD, where a KDYF motif is found in Chondrichthyes and a KEFF motif in all other bony fish and tetrapods. Certain motifs are unique to the specific CtBP paralogs, and are completely conserved across species; these include the very C-terminal sequences, which were also found to be highly conserved in protostomes (green highlight; **Figure 7B**). It is likely that these distinct motifs represent variations that arose relatively soon after CtBP gene duplication. Examples of motifs that are common to CtBP1 family paralogs include central VEGIV motifs, that are clearly related to, but distinct from, the somewhat less conserved CtBP2 MEGMV motif (pink highlight; **Figure 7B**). More ancient motifs such as PELNGA, appearing just N-terminal to deeply conserved aromatic residues (W) of the central block, appear to have been present in the last common ancestor of vertebrates and echinoderms (blue highlight; **Figure 7B, S7C**).

### 9) Conservation of CtBP paralogs across Sarcopterygii

An examination of CtBP forms in these vertebrate species reveals very different levels of evolutionary variation. The super class Sarcopterygii comprises the more basal lungfish (Dipnoi) and coelacanth (Coelacanthimorpha), as well as more recently derived tetrapods including mammals, birds, reptiles, and amphibians (**Figure 10A**). The sequences of both the CtBP1 and CtBP2 CTDs are very highly conserved (**Figure 10B, C, S8A, B**). We find evidence of some substitutions at specific sites, with the length and sequence having high conservation across all Sarcopterygii sampled, and much more within each particular class. Intriguingly, we found that the CtBP2 of amphibians diversified in the length and sequence (**Figure 10C, S8B**). Conserved truncated versions of the CTD observed in some amphibians terminate immediately C-terminal to the central block motif, suggesting that the first portion of the CTD, which includes highly conserved aromatic/hydrophobic residues, may possess a function that is conserved even in these variants (**Figure S8B**). These amphibians are the only Sarcopterygii to have modified the CtBP2 tail, while some Sauria also produce a second short variant (data not shown). In mammals, the only major deviation from the canonical sequence was found in bats (Chiroptera), which are the only order to have short isoforms of both CtBP1 and CtBP2 (**Figure S8D**). The third gene in Sarcopterygii, CtBP1-like, which was lost solely in mammals, has more variation than the other paralogs (**Figure 10D, S8C**). Short variants of CtBP1-like exist in Sauria as well (data not shown). Over the course of ∼400 million years of evolution of Sarcopterygii, we find very high conservation of the CtBP CTD. Although these vertebrates have a longer generation time and slower rate of molecular evolution than insects (which also evolved over 400 million years), the almost invariant nature of the CTD suggests that in Sarcopterygii, conservation of CtBP sequence is critical for function.

**Figure 10.**
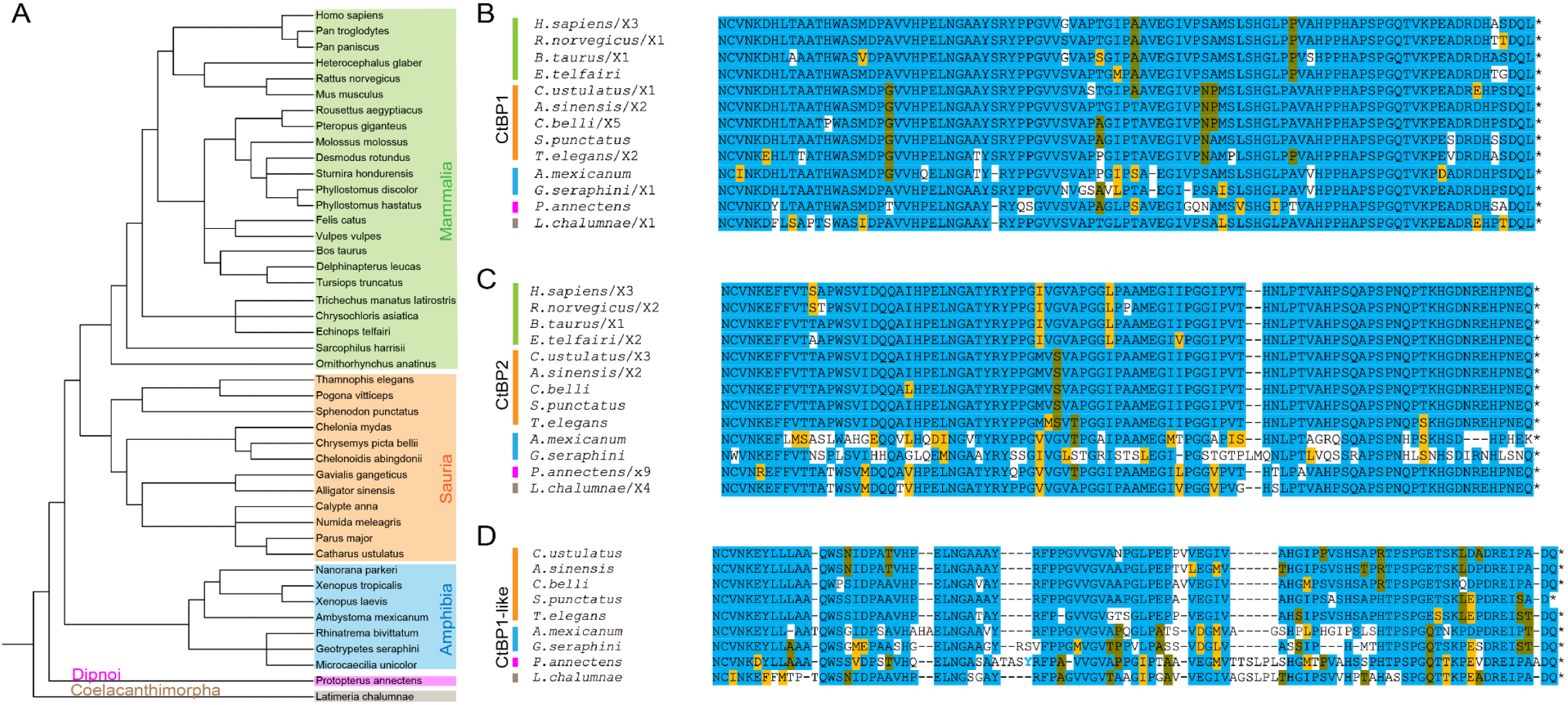
Conservation of CtBP CTD sequences in Sarcopterygii. **A)** Phylogenetic tree of Sarcopterygii species analyzed. Colored boxes indicate major classes of Sarcopterygii, including mammals, sauria, amphibians, lungfish, and coelacanth. Vertical lines in B-D correspond to the groups shown in panel A. **B)** Representative alignment of CtBP1 isoforms indicates that the CtBP1 CTD is highly conserved among selected lobe-finned fishes. A handful of species also express a shorter version of CtBP1 (not shown), while all other species have only a long variant of CtBP1. **C)** Representative alignment of CtBP2 isoforms indicates that the CTD is also highly conserved but is less well-conserved than CtBP1 among selected amphibian species. Some bats (**Figure S8D**) and sauria have short versions of CtBP2. **D)** Representative alignment of CtBP1-like isoforms. CtBP1-like paralogs are absent in mammals.

### 10) Actinopterygii express up to five CtBP genes

Actinopterygii are the second and most speciose branch of the Euteleostomi clade of bony vertebrates. In Actinopterygii, additional CtBP paralogs have arisen, likely through the additional rounds of whole genome duplications documented in fish. In particular, teleosts experienced a third round of genome duplication, termed Teleost-specific Genome Duplication, which dates to 225-333 million years ago (Berthelot *et al*. 2014). We selected sixteen fish that cover all major groups of ray-finned fishes, including many teleost fish and some more basal, ancient fish such as the bichir (*P. senegalus*), paddlefish (*P. spathula*), and gar (*L. oculatus*). Up to five unique CtBP proteins were found in select species including some Teleostei (*A. anguilla*, the European eel; *M. cyprinoides*, the Indo-Pacific tarpon; and *S. formosus*, the Asian arowana) (**Figure 11A, B**). All of these ray-finned fishes express the canonical CtBP1 and CtBP2 paralogs, which differ slightly in their CTDs, but are very highly conserved (**Figure 11C**). We found evidence for short isoforms in some Actinopterygii; the CtBP1-short CTDs are highly conserved, while the CtBP2-short CTD sequences are characterized by a greater degree of variation, even within a single species (**Figure 11D**). Core motifs, such as the NCVNKEY at the beginning of the CTD and the conserved central block, are present in all isoforms analyzed.

**Figure 11.**
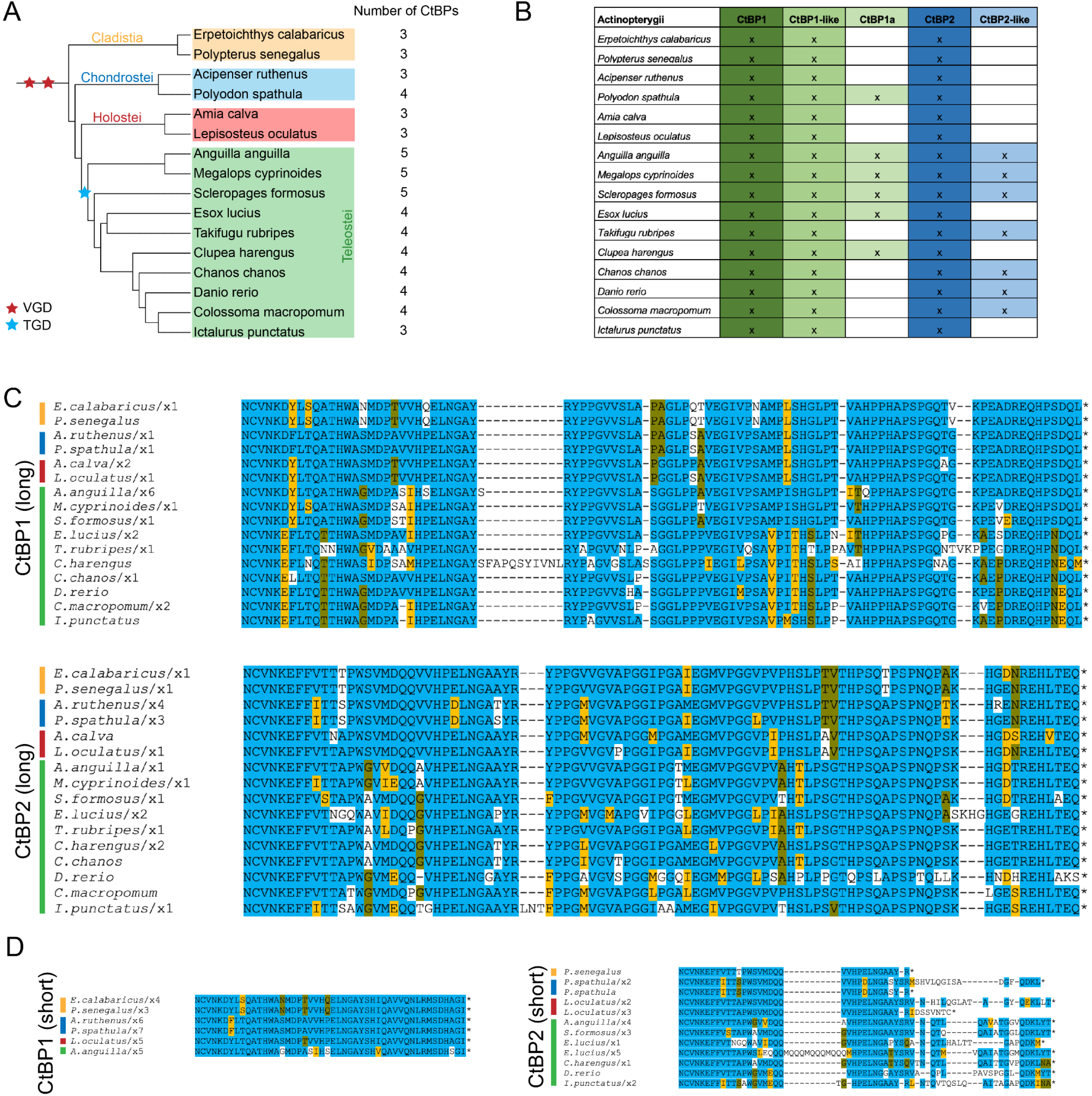
Actinopterygii possess up to five CtBP genes. **A)** Phylogenetic tree of ray-finned fishes divided into four classes/infraclasses. Red stars represent Vertebrate Whole Genome Duplication events (VGD) and the blue star represents a Teleost-specific Genome Duplication event (TGD). **B)** Chart representing CtBP isoforms encoded in each species’ genome. **C)** Alignment of CtBP1-long and CtBP2-long isoforms in the fish indicates that the CTD is well conserved. **D)** Alignment of CtBP1 and CtBP2 short isoforms. Evidence for short isoforms is limited to a subset of species. The CTD sequences are much more conserved in CtBP1-short than in CtBP2-short.

All Actinopterygii have a CtBP1-like paralog, as was observed in most Sarcopterygii, and those with four or five CtBP paralogs express what we have termed CtBP1a and CtBP2-like, determined based on the degree of similarity in the dehydrogenase core to CtBP1 or CtBP2 (see Materials and Methods; **Figure 11B**). Alignments of each of these additional CtBP proteins to one another across Actinopterygii confirms their high degree of conservation and paralog-specific sequences (**Figure S9**). In summary, we find that across 400 million years of evolution of Actinopterygii, gene duplications played a big role in the diversification of this family, while retaining key features of CtBP seen in Sarcopterygii. We hypothesize that the additional CtBP paralogs may function in new roles distinct from transcription; these may include cytoplasmic functions in the Golgi and in synaptic vesicles and neurons, as found for the RIBEYE variant of CtBP2 (Schmitz *et al*. 2000; tom Dieck *et al*. 2005).

### 11) Structural properties of CtBP C-termini

Our survey of the C-termini of CtBP forms across bilaterian diversity indicates that many, but not all, lineages have retained primary structural elements that presumably reflect important functional properties. “Canonical” CTD structures include conserved hydrophobic/aromatic N-terminal residues, an aromatic (Y, F or W) adjacent to the central block, and paralog-specific C-termini with similar arrangements of lysine and acidic residues. This general structure is found across deuterostomes, where most CTDs range in length from 90 to 100 residues (**Figure S10C-E**). In the vertebrates, the CTD of CtBP1 paralogs in most tetrapod lineages have few modifications, and remain 90 to 100 in length (**Figure S10C**), in contrast to the more dynamic length changes evident even within the Drosophila genus. As noted in the analysis of protostome CtBP structure, certain lineages have independently substituted canonical CTD sequences with novel structures, leading to a greater diversity of lengths (**Figure S10A**).

Are there common properties of these diverse CTD sequences, which may reveal common functions? Intrinsically disordered regions (IDR) are found to have characteristic skewed composition of amino acids, with enrichment in structure-disrupting and charged residues, and depletion in hydrophobic (especially aromatic) amino acids (Habchi *et al*. 2014). Considering overall amino acid composition of all CtBP CTDs, the occurrence of hydrophobic residues (M, I, V, L, F, Y, W) is around 21% in Protostomes, while some lineages average closer to 10% (**Figure S10B**). Deuterostomes range from 16 to 27%, averaging ∼25% across the paralogs (**Figure S10F**). This is lower than the frequency found in the CtBP structured dehydrogenase domain, which is around 33% hydrophobic in the fly CtBP and human CtBP1. In comparison to experimentally validated repressor domains in the human proteome, which average around 45% hydrophobic content, the CtBP CTD also has much lower hydrophobicity (A and P residues were included and V excluded, compared to our method; Soto *et al*. 2022).

Many IDRs are polyampholytes, with both positively and negatively charged residues. Strong vs. weak polyampholytes are determined by the proportion of charged residues and their distribution across the peptide, with IDRs that have well-mixed charged residues allowing for repulsion of opposite charges and the adoption of a random coil conformation (Habchi *et al*. 2014; Das and Pappu 2013). For the CtBP CTD, we calculated the proportion of positively charged (K and R) and negatively charged (D and E) residues (**Figure S11A-D**). We find that there is some variability across Protostomia, with Annelids having only 6% of the CTD composed of charged residues, while Nemertea have 18%. Across Deuterostomia, there is much less variability with both CtBP1 and CtBP2 CTDs composed of just under 15% charged residues. Using CIDER (Holehouse *et al*. 2017), we find that based on the low hydrophobicity, and high content of charged residues, these CTDs are considered “weak polyampholytes and weak polyelectrolytes” (FCR <0.3, and NCPR <0.25). These properties suggest that the CTDs may form defined structures in a facultative manner.

IDRs are often enriched in proline, glycine, and alanine residues, which are considered structure-breaking residues (Habchi *et al*. 2014). Proline is the most disorder-promoting of all amino acids, with a disorder propensity score of 1.0 while alanine and glycine have 0.45 and 0.43 respectively (Theillet *et al*. 2013). We analyzed the composition of the primary peptide sequences of CtBP CTDs across Metazoa, calculating total content of proline, glycine, and alanine residues. The composition remains similar across protostomes and deuterostomes, with P, G, and A residues accounting for 35-45% of the entire CTD in most lineages regardless of the CTD length or primary sequence (**Figure S11E-H**). These values are much higher than in the dehydrogenase core, where P, G, and A only make up about 22% of the primary sequence in the fly CtBP and human CtBP1.

We also performed secondary structure predictions to determine whether any CTD species have discernible structures. Using PSIPRED and Robetta (Buchan and Jones 2019; Baek *et al*. 2021) we found that most protostome CtBP CTDs have predicted unstructured domains (**Figure 12A**), with predicted short alpha helices correlated to nonconserved alanine-rich insertions in some species, such as in Drosophila (data not shown). Interestingly, a much greater degree of predicted structure is found in highly derived, lineage-specific CTD sequences in certain mites, nematodes, tardigrades, and flatworms (**Figure 12B**). These possibly much more structured domains, which do not bear structural resemblance, may play specialized roles in these species. Vertebrate CtBP1 and CtBP2 CTDs were predicted to also be highly unstructured, with most having a beta turn or a small alpha helix (**Figure 13**). The predicted beta turns are found within the conserved central block motif, towards the N-terminus of the CTD, and this feature is conserved across representative protostomes and deuterostomes. This structured motif may be important for binding to cofactors.

**Figure 12.**
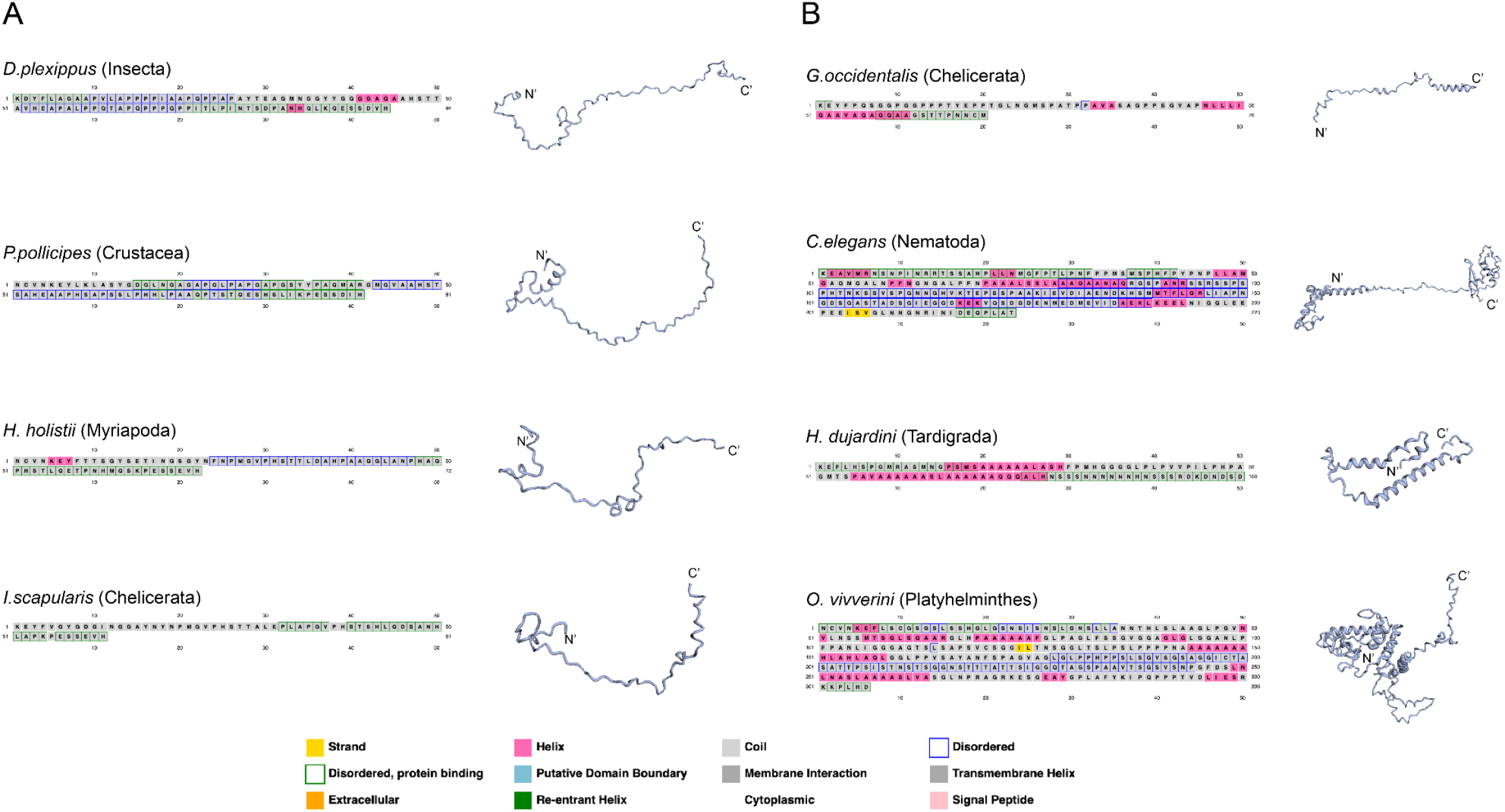
Secondary structure predictions of select protostome CtBP CTDs. PSIPRED (boxed amino acids) and Robetta (structures) predictions. The legend on the bottom indicates the significance of colored boxes. **A)** Representative protostomes with canonical CtBP CTD sequences were selected. In all the predicted structures, the CTD is highly disordered, with a beta turn toward the N-terminus, which maps to the central block motif. **B)** CTD sequences and structures from four species with derived CTDs are shown (specific mites (chelicerates), nematodes, tardigrades, and Platyhelminthes). These secondary structures were found to have less disordered regions and instead had a higher number of alpha helices predicted. These are the only CTDs that are predicted to have a distinct structure. The N- and C-termini are indicated.

**Figure 13.**
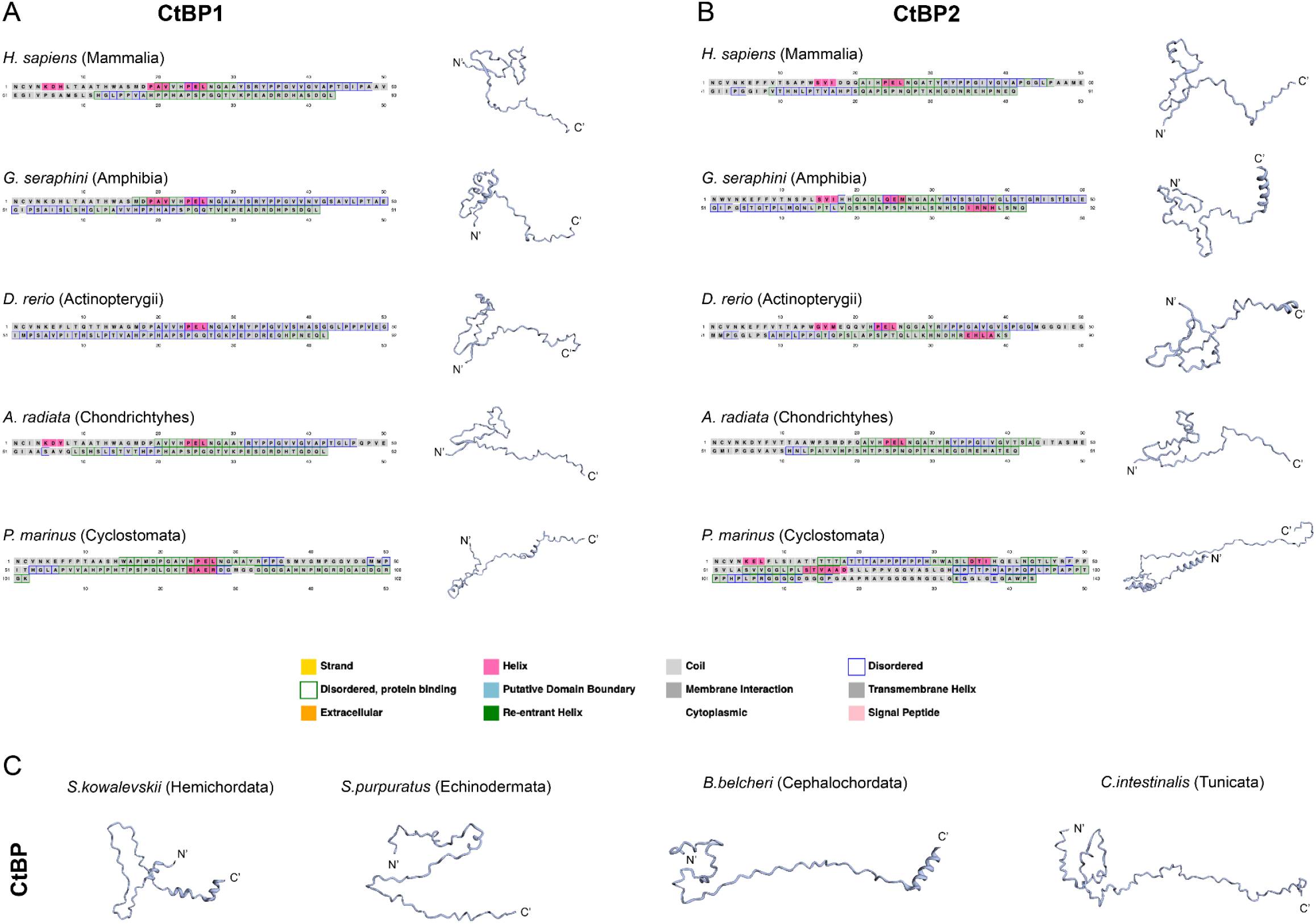
Secondary structure predictions of select deuterostome CtBP CTDs. PSIPRED (boxed amino acids) and Robetta (structures) predictions. The legend on the bottom indicates the significance of colored boxes. **A)** Representative structures for the CtBP1 CTD from Sarcopterygii (Mammalia & Amphibia), Actinopterygii, cartilaginous fish (Chondrichthyes) and jawless vertebrates (Cyclostomata). **B)** Representative structures for the CtBP2 CTD from the same species shown in A. **C)** Predicted secondary structures for non-vertebrate deuterostome CTDs. These CTDs are predicted to be unstructured, with some small alpha helices on the C-terminal portion. The widely conserved central block motif is found to form a beta-turn in most deuterostome species. The N- and C-termini are indicated.

### 12) Diversity of the N-terminal domain of vertebrate CtBP

Aside from the great variation seen in the C-terminal domain across bilaterians, variations in CtBP also exist in the NTD. This is particularly evident in Gnathostomata, the jawed vertebrates, who have diversified CtBP2 through usage of alternative transcriptional start sites to create isoforms with extended NTDs. In mammals, this CtBP2 isoform with an NTD extension is termed RIBEYE, and it produces a protein with a 572 residue extension on the N-terminus (Schmitz *et al*. 2000; **Figure S12A**). RIBEYE has been shown to localize to synaptic vesicles in the retina and in sensory neurons (Schmitz *et al*. 2000; tom Dieck *et al*. 2005). To determine where the RIBEYE isoform first arose outside mammals, and whether non-mammalian vertebrates have a conserved RIBEYE isoform, we analyzed NTD sequences of CtBP2 proteins across Vertebrata.

There is no evidence of long NTD isoforms in the single Cyclostomata species analyzed. Most Gnathostomata express CtBP2 isoforms with extended RIBEYE-like NTDs, with lengths of 550-620 residues (**Figure S12B**). The lack of long NTDs observed in select species may be due to lack of expression in the tissues from which transcriptomic data was collected, or poor detection during sequencing, rather than a loss of long NTD isoforms in these vertebrates. Compared to the human RIBEYE sequence, mammals have 65-80% sequence conservation, while other Sarcopterygii have 45-55% conservation. Actinopterygii and Chondrichthyes also have 40-50% sequence conservation, similar to birds, reptiles, and amphibians (**Figure S12B**). While the levels of conservation among RIBEYE domains is lower than that found in the catalytic core and CTD, there are blocks of sequences that are highly conserved across Gnathostomata (**Figure S12C**). For instance, the MPVPS-like motif at the translational start site is conserved across the sampled gnathostomes, and there are additional 5-10mer motifs that are highly conserved and scattered across the RIBEYE sequence (**Figure S12C**). Species with CtBP2-like isoforms (found only in Teleostei), long NTD isoforms are sometimes present, confirming that CtBP2-like originated from a CtBP2 duplication event. Only a few teleost species have long CtBP2-like NTDs, which are either ∼300 or 700 residues long. Those with 700-residue NTDs (*C. macropomum* and *C. chanos*) have sequences that resemble the human RIBEYE, with about 40% primary sequence conservation, similar to that seen with the CtBP2 of select Actinopterygii (data not shown). We also find conserved 10mer motifs scattered throughout, with insertions of polyQ tracts in the fish CtBP2-like NTD, which results in the longer observed lengths (data not shown). Taken together, these results indicate that the extended CtBP2 N-terminus originated in the last common ancestor of Gnathostomata, and that after gene duplication producing the CtBP2-like paralog in certain fishes, the extended NTD was retained.

### 13) Modifications to the CtBP CTD may add an additional layer of regulation

We have shown that over longer evolutionary times, novel forms of CtBP have developed at the gene level through wholesale adoption of unique CTD sequences, isoform production using alternative splicing, gene duplication, and alternative promoter usage. Variation in CtBP is also found at the protein level, via post-translational modifications (PTMs). A few studies have focused on the functional impact of CtBP PTMs, but most of our knowledge is derived from mass spectrometric analysis of the human proteome from diverse samples, which has uncovered some modifications to the CtBP1 and CtBP2 proteins. Not surprisingly, the CtBP CTD can undergo many PTMs, which is a common feature of IDRs because they are accessible to enzymes for modifications (Musselman and Kutateladze 2021). The CtBP1 CTD is phosphorylated at S422 by HIPK2, which triggers CtBP degradation and cell death, and is sumoylated at K428 by SUMO-1, which allows for its nuclear localization (Zhang *et al*. 2003; Zhang *et al*. 2005; Wang *et al*. 2006; Kagey *et al*. 2003; Lin *et al*. 2003; **Figure S13A**). These residues are completely conserved across vertebrate CtBP1 and CtBP1-like, and also among some non-vertebrate deuterostomes, suggesting that the CtBP1 tail can be modified and regulated in a similar manner in these species (**Figure S13B**). CtBP2 is phosphorylated on residues S365, T414 and S428. HIPK2 phosphorylates S428, but the impact of this and other modifications have not been experimentally determined (Bian *et al*. 2013; Dewi *et al*. 2015; **Figure S13A**). Only T428 is conserved across vertebrates, while the other residues show lower conservation (**Figure S13C**).

To determine whether PTMs such as phosphorylation and sumoylation may involve conserved portions of the CTD in our selected species, we used predictive PTM software. We determined putative sumoylation sites using JASSA v4 (Beauclair *et al*. 2015). We find that many invertebrates with a canonical CTD sequence including insects, chelicerates, and some Spiralia, have high consensus SUMO motifs, usually in the extreme C-terminus. Many of the derived CTD sequences from Nematoda also have a predicted sumoylation site. The majority of Sarcopterygii CtBP1 sequences have a strong SUMO consensus motif in the extreme C-terminus, while Actinopterygii have a weak motif. The CtBP2 CTDs lack SUMO motifs, suggesting a different form of regulation and activity. We also predicted possible phosphorylation sites in the CTDs, as IDRs have been shown to be particularly enriched in phosphorylated residues (Habachi *et al*. 2014). Using NetPhos 3.1 (Blom *et al*. 1999), we find that the Y and S/T residues of the vertebrate central block are predicted phosphorylation sites, as are the same residues in the invertebrate central block motif. The high conservation of this motif across Bilateria, and its predicted phosphorylation status may point to an important role in regulation of CtBP activity. Additionally, protostome-specific motifs (AHSTTP and the terminal SEVH ending) are also predicted phosphorylation sites, again pointing to positive selection perhaps due to an important regulatory role.

### 14) Evolutionary variation in the conserved dehydrogenase core

The well-structured dehydrogenase domain of CtBP shows much higher sequence conservation than the CTD, and across longer evolutionary time. However, some variation in the core is found between species; in Diptera, a number of species generate alternative splice forms that affect the VFQ tripeptide motif which is predicted to be in an unstructured loop on the surface of the protein (**Figure S14A**; for *D. melanogaster*, see **Figure 1C**). VFQ is present across most insects, with some variations, and is found in some arthropods, such as in Crustacea (IFQ) and Myriapoda (IHQ), but the motif is not conserved across Protostomia (**Figure S14A**). It presumably is only spliced out in Diptera, as there is no evidence that there are isoforms without VFQ in other insects or protostomes. The effect of its retention or removal is not yet known, but due to the conservation of its alternative splicing, we suspect it plays a role in the structure of the protein or the recruitment of cofactors. Additional core variations are found more broadly in arthropods, such as a five residue insertion in some splicing isoforms of select insects and chelicerates, N-terminal to the start of the CTD (**Figure S14A**). Interestingly, this motif maps just C-terminal to the VFQ, also in a predicted unstructured portion of the protein on the surface of the structure, and away from the tetramerization interface. Among the protostomes, there are several spiralian and crustacean species that have 1-15 amino acid motifs that are inserted or deleted, which are unique to only those species, and presumably arose much later in their evolution since they are not alternatively spliced in other species.

Among the deuterostomes, the only conserved alternatively spliced motif is SF, found ∼50 residues before the start of the CtBP1 CTD (**Figure S14B**). SF is alternatively spliced in select Actinopterygii and Sarcopterygii, but not in all examined species. Interestingly, this motif also maps to an unstructured loop in the human CtBP1 protein, and overlaps the dipteran VFQ motif, suggesting that its alternative splicing is significant, either because it was retained, or independently arose in these two lineages. Other vertebrates have variations in their core that are species-specific, and may be transcripts that are expressed at only particular developmental time points or tissue types. Most of these variations in the core are found on the surface of the protein, in regions that lack secondary structure (data not shown). Additionally, they are far from the tetrameric interface, which suggests that tetramerization may not be impacted by these variations in the core, but perhaps interactions with cofactors are altered.

The catalytic triad, which is emblematic of CtBP as an ancient dehydrogenase, is conserved in all bilaterians, aside from nematodes, which have lost one of the three (**Figure S1**). Interestingly, all metazoans retain these residues, but the *Arabidopsis thaliana* ANGUSTIFOLIA homolog does not, consistent with the divergent function of the plant protein in the cytoplasm. Tetramerization residues found in the core, which have recently been shown to be necessary for CtBP2’s activity as a transcriptional repressor, are also highly conserved. In the human CtBP2, five residues were shown to be necessary for tetramerization, which allows for transcriptional repression: S128, A129, R190, G216, and L221 (Bellesis *et al*. 2021). Between the human CtBP1 and CtBP2, four of these are conserved (not S128; Raicu *et al*. 2021). In bilaterians, the R, G, and L residues are completely conserved, and SA is either GY, GF, or GV. Perhaps tetramerization is a more broadly conserved structural feature of these proteins.

## Discussion

Our comparative phylogenetic study demonstrates that CtBP is a bilaterian innovation, with virtually all orthologs possessing an unstructured C-terminus, usually of about 100 residues. Although initial observations of CtBP protein sequences suggested that the CTD was not conserved, here we demonstrate striking patterns of deep conservation (Kim *et al*. 2002; Nicholas *et al*. 2008). Across Metazoa, the CTD is highly conserved in length, in its propensity for disorder, and in certain blocks of sequence that are found in most species. The long C-terminus is found in virtually all lineages, with additional shorter alternative isoforms arising through alternative splicing independently in a number of insects and vertebrates. Interestingly, there are lineages where the sequence and structure of the C-terminus has independently undergone radical transformations; in mites and tardigrades, the length is maintained but the sequence has diverged, while divergent flatworm and nematode CTD sequences extend to several hundred residues. In vertebrates, additional diversification of CtBP is found through gene duplication, with up to five unique genes encoded in certain fishes. Diversification of the CtBP CTD may have implications in gene regulatory networks, and more broadly in evolutionary transformations of bilaterians.

Viewed broadly, this analysis of CtBP evolution shows some parallels to previous studies of other components of the bilaterian transcriptional machinery, while raising some still unanswered questions. From pioneering work by Lewis and others, reverse engineering transcriptional systems has uncovered important cis and trans variations in components of the transcriptional machinery that drive profound evolutionary transformations in the metazoan body plan (Lewis 1978) Those variations affecting DNA-binding transcription factors, such as Hox proteins, provide some of the best-known cases (Pearson *et al*. 2005). On the other hand, the potential impact on morphological evolution stemming from variation in the core regulatory machinery that is responsible for expression of most genes, is less well known. Indeed, initial biochemical studies of the basal transcriptional machinery, including RNA polymerase II and associated factors, emphasized the conservation of a largely invariant and nearly universal collection of components specific to eukaryotes, underlining the early emergence of these factors in the last common ancestor. However, more recent work has demonstrated lineage-specific features of this machinery, including the diversity of factors within the TFIID complex (TBP and TAFs) pointing to the specialization of even the pleiotropic core machinery (Li *et al*. 2009; Goodrich and Tjian 2010).

As we document for CtBP, a significant source of variation within the core transcriptional machinery is found in intrinsically disordered regions (IDRs). Overall, IDRs feature low sequence complexity, low hydrophobicity, and are heterogeneous in their conformation (Shukla *et al*. 2022). They can self-associate, adopt structured conformations in association with cofactors, or participate in flexible interaction surfaces (so-called “fuzzy” complexes) (Shukla *et al*. 2022). These properties appear to lend IDRs a particularly active role in evolutionary change, as they can tolerate substitutions and still perform their diverse functions (Musselman and Kutateladze 2021; Pajkos *et al*. 2021; Shukla *et al*. 2022). What functions might be associated with the CtBP CTD? Studies of a number of IDR-containing proteins point to a diversity of roles, including roles in regulation of DNA binding, cofactor recruitment, anchors for post translational modifications, homodimerization, and adopting defined structures in a larger complex. It is notable that structural studies of CtBP that have emphasized its unstructured CTD used purified protein, thus it is possible that the CTD is highly structured when combined into a complex of other interacting partner proteins.

We think that the most attractive model for the CTD of CtBP is provided by a transcriptional cofactor that is derived from another class of hydroxyacid dehydrogenases, N-PAC (also known as GLYR1). This protein, which like CtBP can form homotetramers, possesses a disordered N-terminus. The IDR associates with the LSD2 demethylase, and a portion of its sequence adopts a defined structure to assist LSD2 to access histone tails (Marabelli *et al*. 2019). The long flexible N-terminal region has been suggested to allow the tetrameric complex to simultaneously span several nucleosomes, coordinating the action of this chromatin modifying complex. Similarly, those conserved portions of CtBP’s CTD may form defined structures in the context of a larger complex, while also contributing to regulation via posttranslational modifications and possible long-range interactions. Ongoing advances in structural biology will likely deliver important information on such multiprotein complexes, which will generate important hypotheses relating to CtBP, such as the expected impact of CtBP-short forms lacking a CTD. How the lineage-specific variations impact gene expression remains a significant challenge that will require an integrated genomic and molecular genetic approach.

## Materials and Methods

### cDNA and peptide sequences

cDNA and peptide sequences for *D. melanogaster* CtBP isoforms were downloaded from flybase (http://flybase.org/; version FB2020_03 and FB2020_05; dm6; Gramates *et al*. 2022). *D. melanogaster* sequences were used as a reference to retrieve cDNA and peptide sequences using NCBI blastn and blastp (https://blast.ncbi.nlm.nih.gov/Blast.cgi) for human CtBP1 and CtBP2, and most protostomes. The human CtBP1 and CtBP2 sequences were used as a reference to retrieve cDNA and peptide sequences for most deuterostomes. When peptide sequences were not available through NCBI, we translated the available cDNA sequences using the “Show translation” tool on bioinformatics.org (https://www.bioinformatics.org/sms/show_trans.html) selecting the translations for “reading frames 1 to 3”. The open reading frame for *D. melanogaster* and *H*.*sapiens* CtBP sequences were used to determine the correct reading frames for other species. Most of the downloaded sequences were annotated as CtBP or CtBP-like in NCBI; sequences labeled as “dehydrogenase” with <40% similarly were not included in this analysis. Platyhelminthes sequences were retrieved by selecting “Transcriptome Shotgun Assembly” on NCBI BLAST. *Adineta vaga* sequences were obtained from GENOSCOPE Adineta vaga genome browser (https://www.genoscope.cns.fr/adineta/cgi-bin/gbrowse/adineta/), *Strigamia maritima* from e!EnsemblMetazoa, *Protopterus annectens* from Marco Gerdol (Biscotti *et al*. 2016), *Euperipatoides rowelli* from https://datadryad.org/stash/dataset/doi:10.5061/dryad.bk3j9kdc0 and *Gyrodactylus salaris* from Paps and Holland 2018. All species used, their taxonomic ID, and genome version are listed in Supplementary File 1.

### Multiple sequence alignments

Peptide sequences were aligned using the MAFFT multiple sequence alignment (https://www.ebi.ac.uk/Tools/msa/mafft/) using the ClustalW output format. All isoforms for a given species were aligned against one another to note differences between isoforms from the same species. A representative isoform from each species was included in the figures. Amino acids that were conserved in >50% of the species in an alignment were colored blue. Chemically conserved amino acids in the same position were colored orange, using the following conservation scheme: Hydrophobic aliphatic amino acids: M, V, I, L; hydrophobic aromatic amino acids: W, Y, F; acidic amino acids: D, E; basic amino acids: K, R; hydroxyl containing amino acids: S, T. Where there was conservation of a second amino acid but in only 25-50% of species, they were colored army green. Amino acids that were not chemically similar and were not conserved across many species in the alignment were left uncolored.

### Phylogenetic trees

We used NCBI to collect taxonomic IDs for all species used in this study. Tax IDs were inputted into phyloT (https://phylot.biobyte.de/) for generation of phylogenetic trees. Timetree (http://www.timetree.org/) was used to compare phylogenetic trees and determine estimated time of divergence between select species.

### Analysis of CtBP CTD properties

We collected the CTD peptide sequences from >200 metazoan species, and used a script (Supplementary file 2) to determine the following for each CTD sequence: length (in Amino Acids, AA), proportion of A, P, G residues, percent hydrophobic residues (M, V, I, L, W, Y, F), and percent charged residues (K, R for positive and D, E for negative). For each property, a species’ longest CTD was used, and properties were averaged by groups (i.e all the insects were averaged together, using a single CTD sequence from each of the selected species). Short CTDs were not used in the analysis aside from the leeches.

### Secondary RNA and protein structure predictions

Secondary structure predictions of RNA were made using RNAstructure (version 6.2) from the Matthews lab at University of Rochester Medical Center (Xu and Matthews 2016). Data was inputted using the Predict a Secondary Structure Web Server with default parameters, with temperature set to 293K. The structure with the highest probability was used. Secondary structure predictions of CtBP CTD peptides were made using PSIPRED Workbench V3.2 (Buchan and Jones 2019). For data input, “sequence data” was selected under the “Select input data type” heading. PSIPRED 4.0 and DISOPRED3 were selected under the “Choose prediction methods” heading. Under “Submission details”, the FASTA peptide sequence of interest was inputted and submitted. Secondary structure predictions of CtBP CTDs were also made into homology models using Robetta from the Baker lab at the University of Washington Institute for Protein Design (Baek *et al*. 2021). For data input, “Submit” was selected under the “Structure Prediction” heading. Under “Protein Sequence”, the FASTA peptide sequence of interest was pasted. RoseTTAFold was used to create models.

### Determination of CtBP paralogs in vertebrates

Several CtBP sequences from the vertebrates with more than two CtBP paralogs were misannotated in NCBI. We performed pairwise sequence alignments and determined the correct CtBP1-like, 1a, and 2-like sequences based on percent conservation to the dehydrogenase core of *H. sapiens* CtBP1 and CtBP2, and to a species’ own CtBP1 and CtBP2. We also determined motifs in the CTD which were representative of CtBP1 or CtBP2 to accurately assign 1-like and 2-like names to the additional proteins. CtBP1-like and 1a sequences have higher conservation in the core and CTD to CtBP1, and the same for CtBP2-like with CtBP2. CtBP1-like and 1a are very similar to each other, but are not 100% conserved in the core within the same species. CtBP1-like/1a CTDs typically start and end with KEYL…PADQ. CtBP2-like CTDs typically start and end with KEFF…LTEQ. We additionally determined whether the cDNA sequences originate from different genomic locations, and found that for the Actinopterygii with up to 5 different genes (i.e *A. anguilla*), they originate from different chromosomes, indicating that they are 5 unique genes. Species where only one CtBP exists but was annotated with a variant name was re-assigned as CtBP (i.e the non-vertebrate deuterostomes with a single CtBP). The two *P. marinus* paralogs were also renamed to CtBP and CtBP-like based on alignments to each other, and to other vertebrate sequences, as described in the text.

### Classification of IDRs

CIDER (Classification of Intrinsically Disordered Ensemble Regions; http://pappulab.wustl.edu/CIDER/analysis/) was used to determine FCR (fraction of charged residues), NCPR (net charge per residue), and the Das-Pappu phase diagram position for representative CtBP CTDs from bilaterian species. CTD sequences were inputted in FASTA format.

### PTM predictions

To predict putative SUMO motifs in the CtBP CTDs, we used JASSA v4 (Joined Advanced SUMOylation site and SIM analyzer; http://www.jassa.fr/; Beauclair *et al*. 2015), which predicts sumoylated lysines based on the presence of a ѱKxα motif, or a variation of it (ѱ = hydrophobic residue; x = any amino acid, α=D or E). We inputted sequences from representative species across Bilateria in FASTA format, with the set parameters. To predict putative phosphorylated S, T, and Y residues, we used NetPhos - 3.1 (https://services.healthtech.dtu.dk/service.php?NetPhos-3.1; Blom *et al*. 1999) by inputting sequences from representative species across Bilateria in FASTA format.

## Acknowledgements

This work was supported by the National Institute for General Medical Sciences (grant number *R01GM124137* to D.N.A), the National Institute of Child Health and Development (grant number *F31HD105410* to A.M.R), the MSU Biochemistry and Molecular Biology Undergraduate Research Fellowship to M.N, the Wielenga Research Fellowship to M.N, the MSU BEACON Luminaries grant to Y.Y, the Moore-Billman Scholarship Award in BMB to K.M.B, and the College of Wooster APEX Fellowship to K.B. We would like to thank Flybase, the researchers that supplied their genomic and transcriptomic data to NCBI, Diego Granados-Villanueva for his assistance with writing the script for the CTD analysis, and Andrew Thompson and Sandhya Payankaulam for helpful discussions.

