## Supplementary Figures for "The cynosure of CtBP: evolution of a bilaterian transcriptional corepressor"

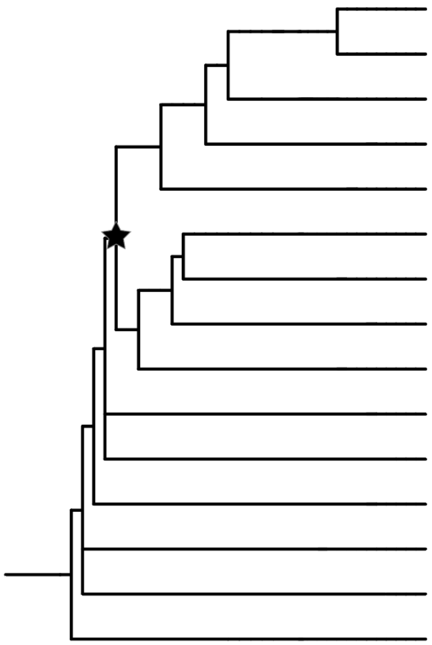

| Group | Representative species' name | % conservation in core | Long CTD | Conserved catalytic triad |
|---|---|---|---|---|
| Sarcopterygii | *H. sapiens* | 100 | X | X |
| Sarcopterygii | *M. musculus* | 99 | X | X |
| Actinopterygii | *D. rerio* | 89 | X | X |
| Chondrichthyes | *A. radiata* | 96 | X | X |
| Echinodermata | *P. miniata* | 80 | X | X |
| Insecta | *D. melanogaster* | 75 | X | X |
| Crustacea | *P. pollicipies* | 72 | X | X |
| Myriapoda | *H. holstii* | 66 | X | X |
| Nematoda | *C. elegans* | 56 | X | |
| Placozoa | *T. adhaerens* | 30 | X | X |
| Cnidaria | *N. vectensis* | <30 | | X |
| Porifera | *A. queenslandica* | 40 | | X |
| Fungi | *E. dermatitidis* | 32 | | X |
| Choanoflagellata | *M. brevicollis* | <30 | | X |
| Viridiplantae | *A. thaliana* | <30 | | |

**Figure S1. Conservation of CtBP features in metazoans and other eukaryotes.** Representative sequences from vertebrates, invertebrates, and non-metazoan eukaryotes were aligned to the human CtBP1 sequence. Percent conservation in the core was calculated as the proportion of conserved residues between the region flanked by the NTD and CTD in humans (everything including and between the RPLVALL and NCVN motifs). We indicate (X) the presence of a long CTD (>80 residues), and the catalytic triad (REH). Bilaterians, indicated by the black star, have canonical CtBP sequences, resembling the known human co-repressor. No CtBP homolog was identified in a representative Ctenophore (*M. leydi*, not shown).

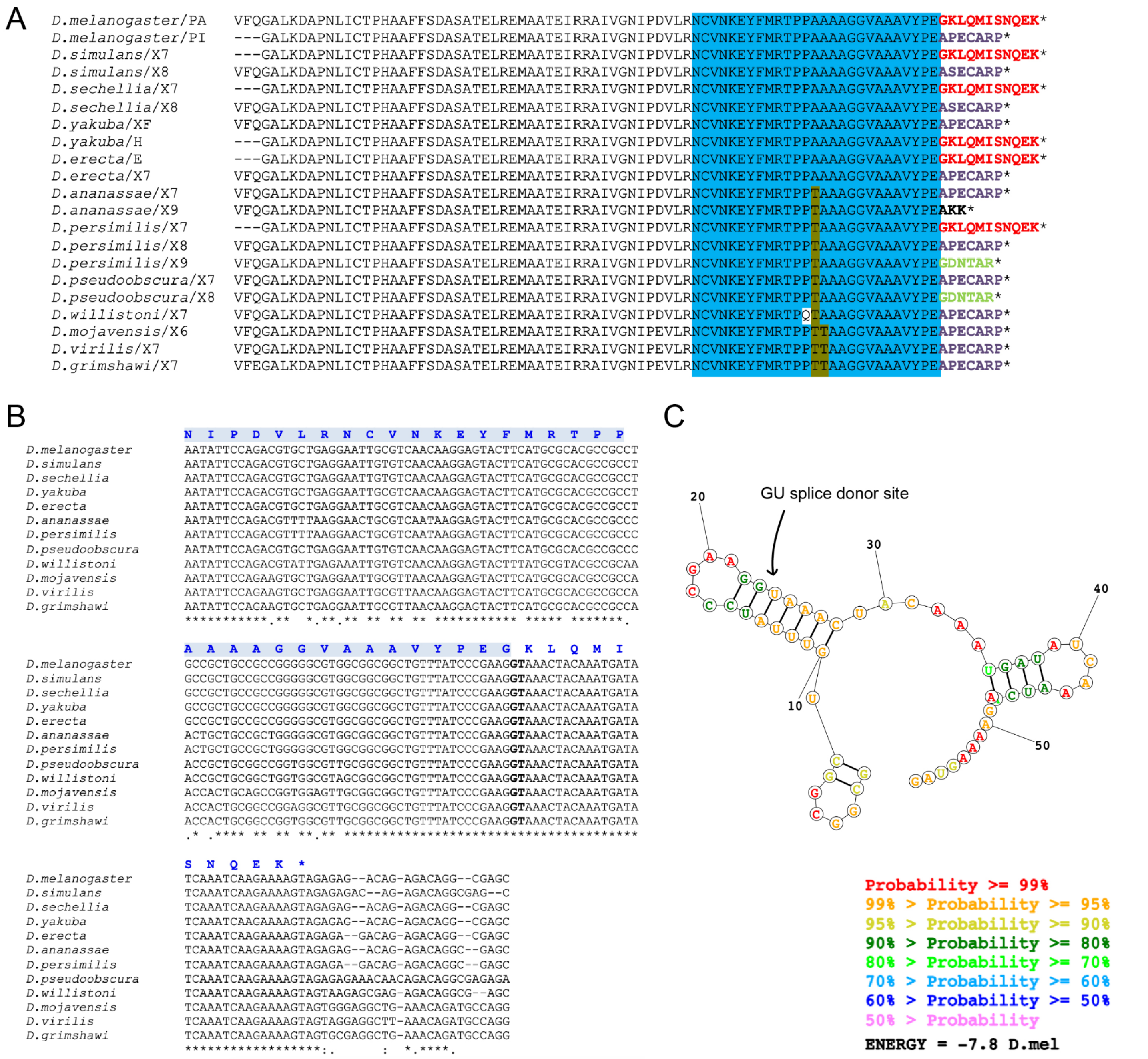


**Figure S2. Formation of $CtBP_S$ through predicted RNA hairpin structures. A)** Alignment of $CtBP_S$ from 12 Drosophila species. SNQEK-like endings are reported for half of the species, while APECARP-like are found in all. Each CTD variant is shown in a different color for the terminal residues. **B)** Alignment of the 3' end of protein coding exon 5 (light blue highlight), and 5' end of adjacent intron among the 12 Drosophila species. The single letter amino acid code indicates the corresponding peptide sequence for *D. melanogaster* for the SNQEK short isoform. This isoform is created through reading through the GT splice donor site (bold) and continuing until a STOP (TAG) codon is reached. We observe 100% conservation (asterisks below alignment) of ~50 nucleotides around the region where splicing normally creates the $CtBP_L$ isoform through use of the donor site. **C)** RNA structure prediction of the region starting with the triple alanine codons (panel B) highlights formation of a possible hairpin structure incorporating the GU splice donor site. Predicted probability of the structure is indicated in the bottom right legend.

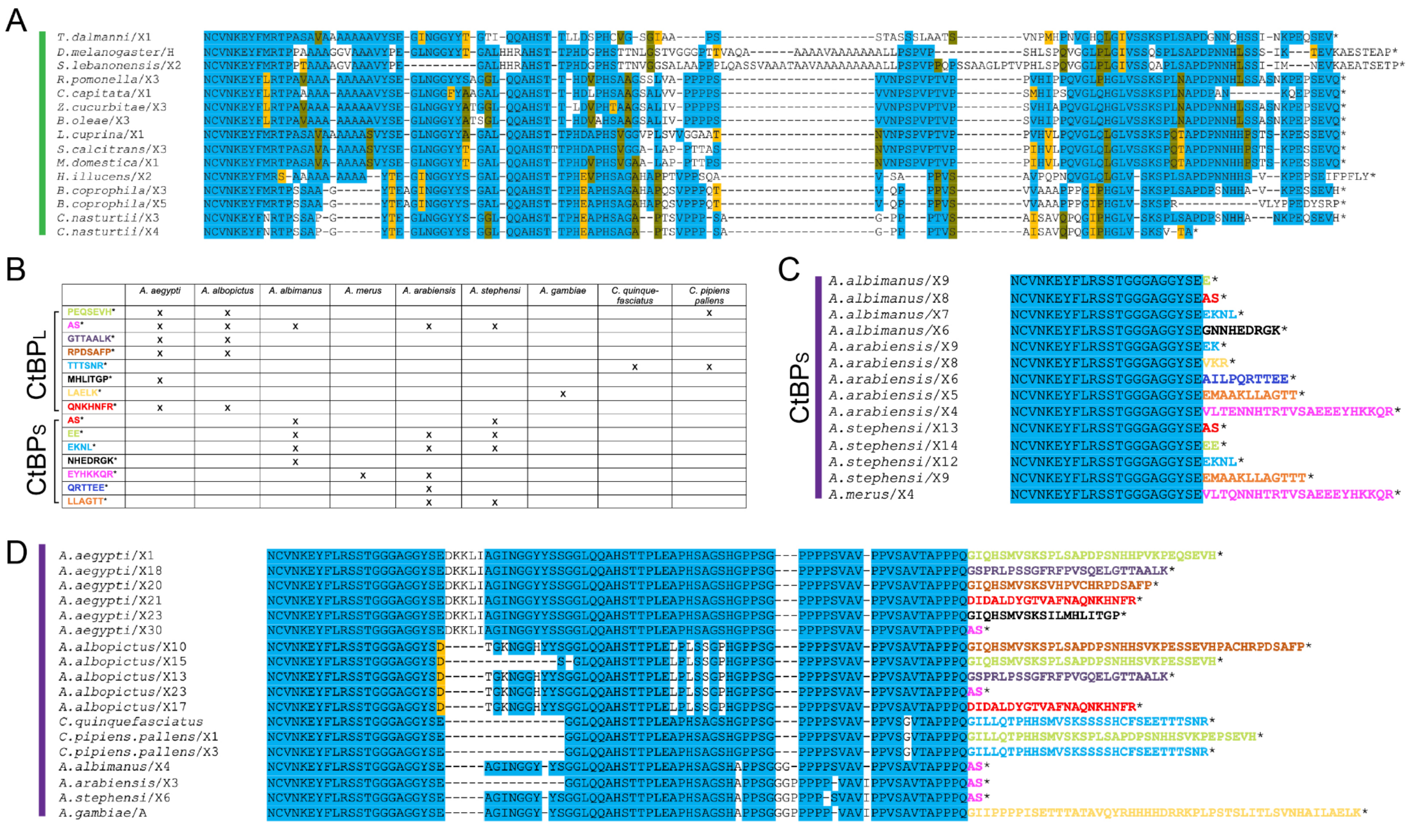


**Figure S3. Lower Diptera have unique CTDs with both long and short isoforms. A)** Alignment of long isoforms from all higher Diptera examined. For all alignments, higher Diptera are indicated by the vertical green line on the left, and lower Diptera by the vertical purple line. **B)** Chart of the long and short CTD variants found in the selected mosquitos (variants are indicated by the very C-terminal sequences e.g. PEQSEVH). All species except *A. merus* have one or more long CTDs, and some species also have a diversity of short CTDs. **C)** Alignment of short isoforms from lower Diptera. Short isoforms are only found in the Anopheles genus, and some species have up to five variants. Variants that are found in more than one species are colored the same (i.e the variant ending in KKQR is found in both *A. arabiensis* and *A. merus*, colored in pink.) **D)** Alignment of long isoforms from all lower Diptera examined indicates that there is high conservation until the terminal PPQ, after which many variations exist across the genera.

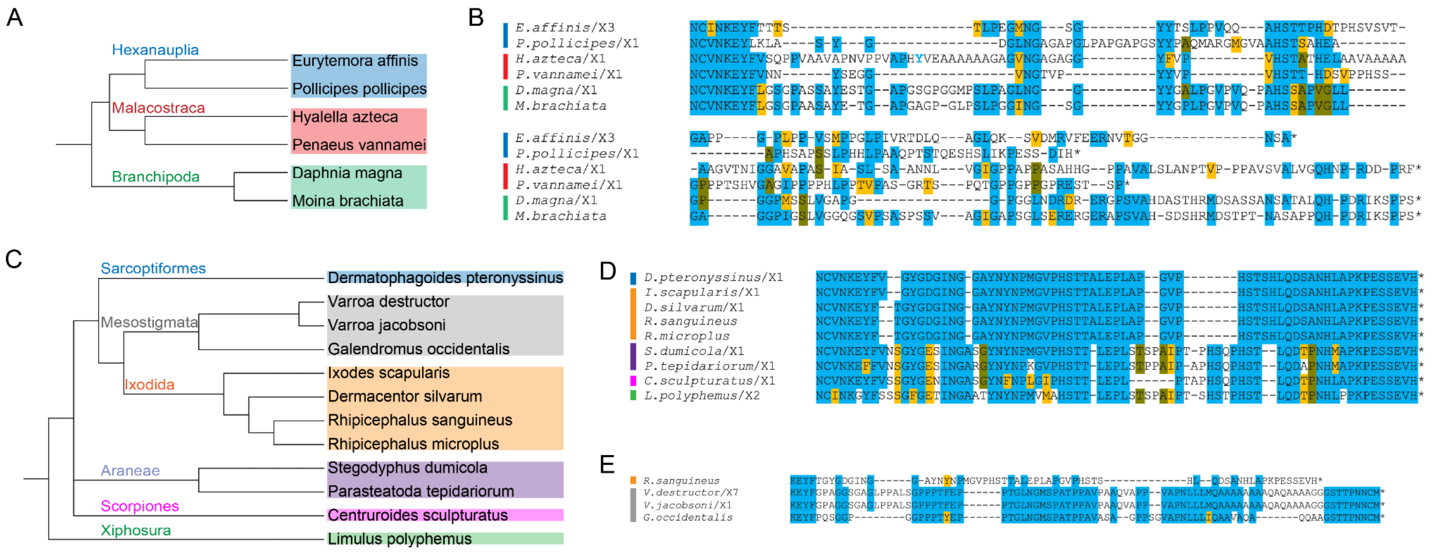


**Figure S4. Alignment of crustacean and chelicerate CtBP CTDs. A)** Phylogenetic tree of three Crustacean groups. **B)** Alignment of six crustaceans indicates that this subphylum experienced diversification particularly affecting the C-terminal sequences after the central block, while some species (*P. pollicipes*) retain presumed ancestral C-terminal sequences (e.g. SDIH). Vertical lines represent species shown in panel A. The tyrosine in light blue (*H. azteca*) is a conserved aromatic residue spaced differently than the other species, but conserved between the NCVN and GLNG--YY motifs. **C)** Phylogenetic tree of select chelicerates including mites, ticks, spiders, scorpions, and horseshoe crab. **D)** Alignment of chelicerate CTDs indicates very high conservation ending with the ancestral SEVH. Vertical bars on left follow the labels in panel C. **E)** Alignment of divergent CTDs from three Mesostigmata mites with the *R. sanguineus* tick, which represents an ancestral sequence.

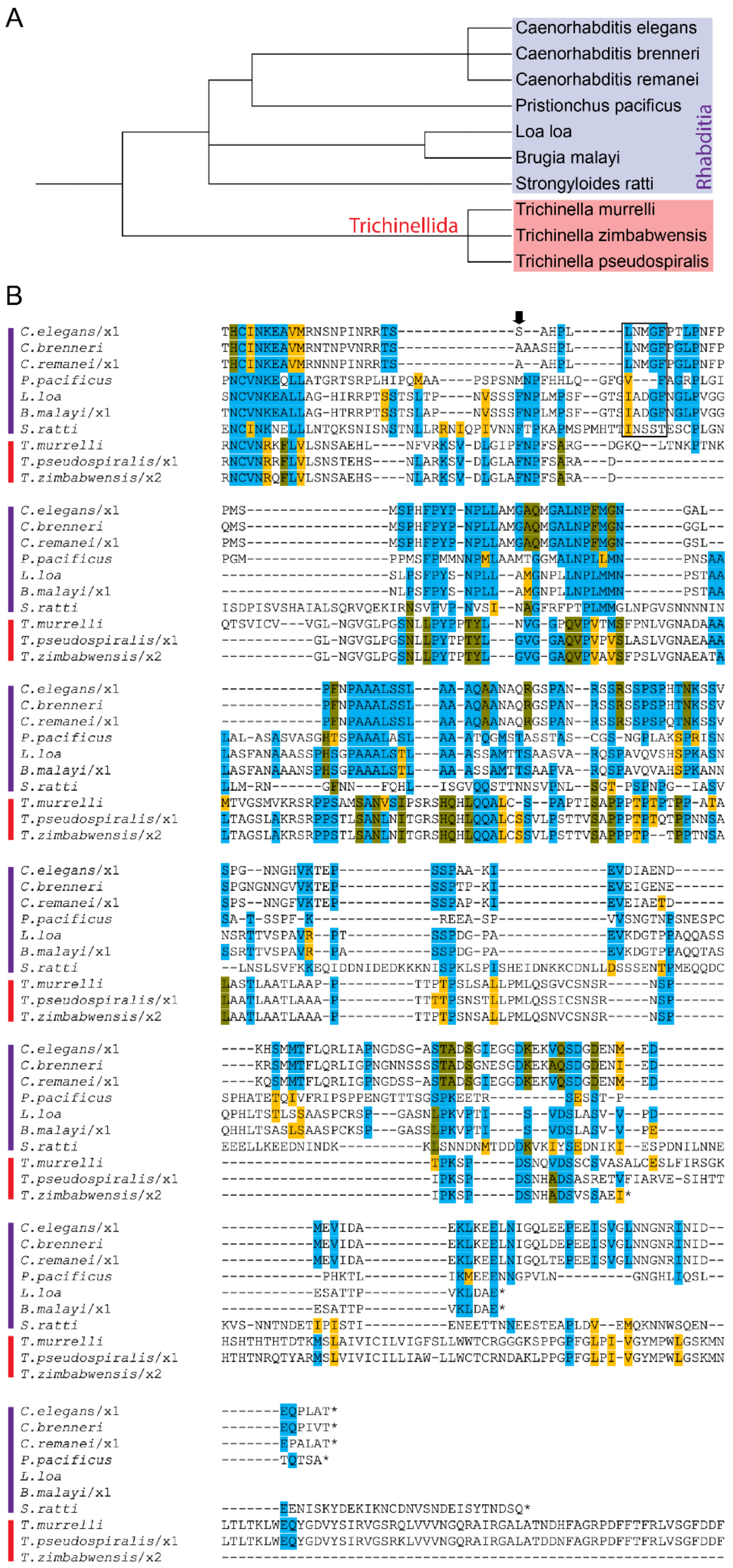


**Figure S5. Nematodes have lineage-specific derived CTD sequences A)** Phylogenetic tree of select roundworms. **B)** Alignment of CtBP CTDs from nematode species covering several genera. Within the same genera (for instance, Caenorhabditis), many features are conserved, but from one genus to the next, there are few similarities in primary sequence. Only portions of the *T. murrelli* and *T. pseudospiralis* sequences are shown; an extension of an additional ~400 amino acids is predicted to complete these CTDs.

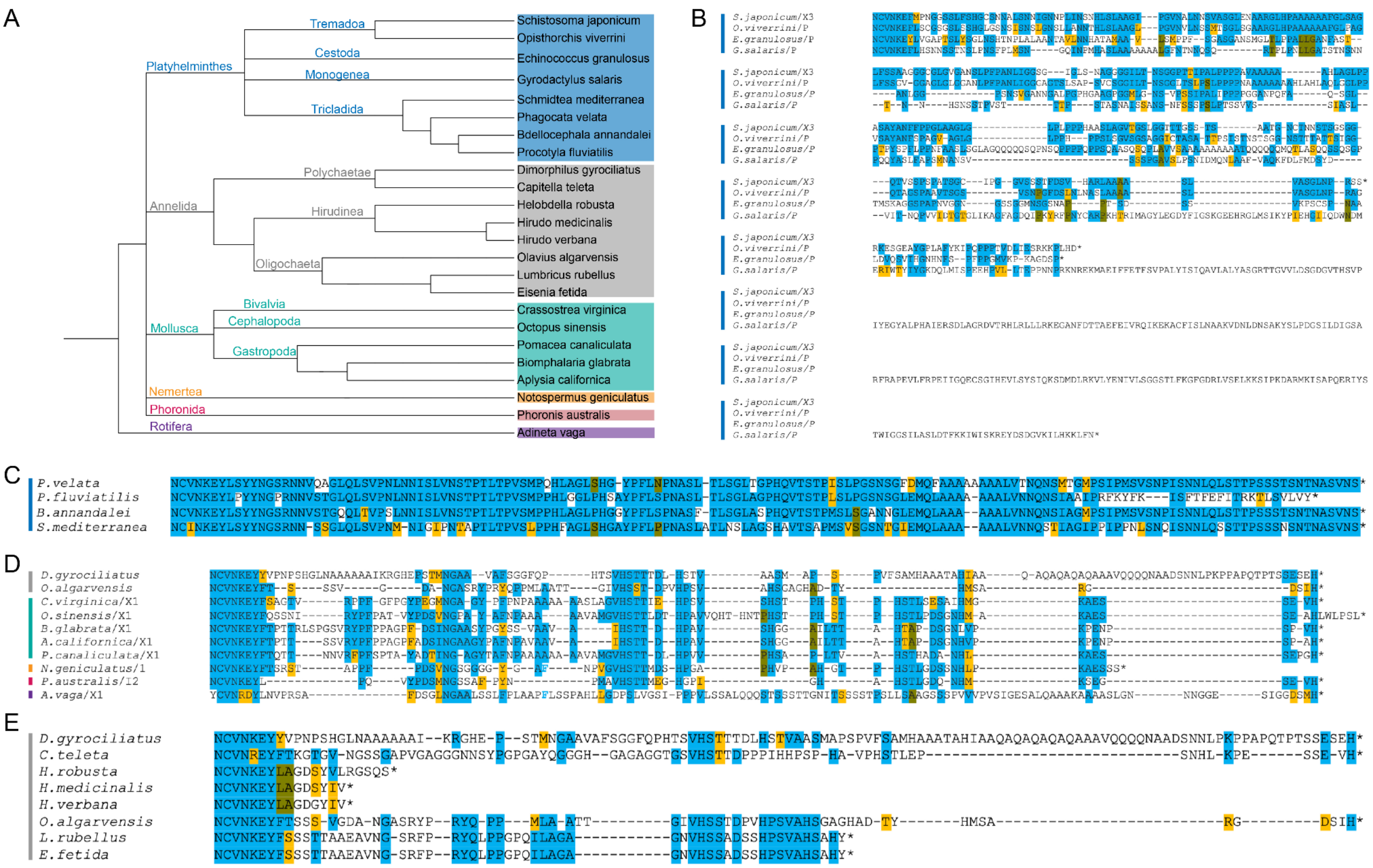


**Figure S6. Spiralia are diverse in their CTDs. A)** Phylogenetic tree of diverse Spiralia species encompassing six phyla. Colored vertical bars in B-E correspond to phyla indicated in A. **B)** Alignment of CTD sequences from trematodes, cestodes, and monogeneids indicates some similarities among these groups, but the sequences are distinct from those found in Triclad Platyhelminthes and the other spiralian species. **C)** Alignment of CTD from four flatworms (Tricladida) showing strong conservation of unique, derived CTD sequences, unlike those found in other spiralians. **D)** Alignment of species within five phyla illustrates the conservation of distinct CtBP CTD motifs, similar to conserved ecdysozoan CTDs. Conserved aromatic (F) residues that are spaced differently in the central block are indicated in blue font. **E)** Alignment of diverse annelids, including earthworms and leeches. The leeches (Hirudinea) are the only species found to express only $CtBP_S$. The Oligochaeta annelids have truncated CTDs that still resemble some of the protostome motifs, such as the central block (VNGSRY/F motif before the highlighted ILAGA).

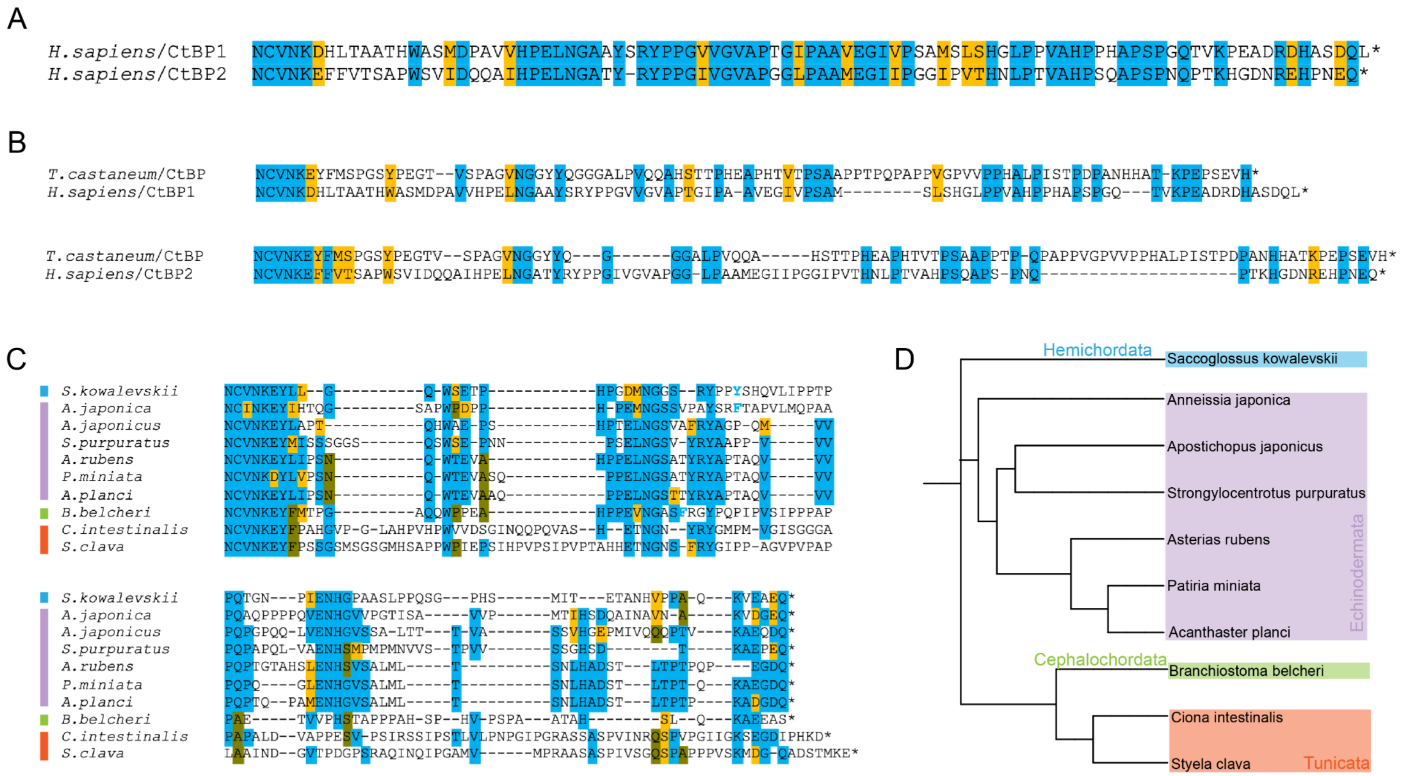


**Figure S7. Evidence for homology among deuterostome CTD sequences. A)** Human CtBP1 vs. CtBP2 CTD alignment. This portion is 50% conserved. **B)** Representative protostome (*T. castaneum* beetle) compared to human CtBP1 CTD and CtBP2 CTD. Some motifs are conserved in protostomes and deuterostomes, such as the central block. **C)** Alignment of non-vertebrate deuterostome CtBP CTD sequences. These species have a single CtBP protein. Aromatic-containing central block sequences are conserved, as are terminal amino acids. Conserved Y and F residues that are spaced differently in the central block are indicated in blue font. **D)** Phylogenetic tree of the four (sub)phyla in Deuterostomia that harbor a single CtBP gene.

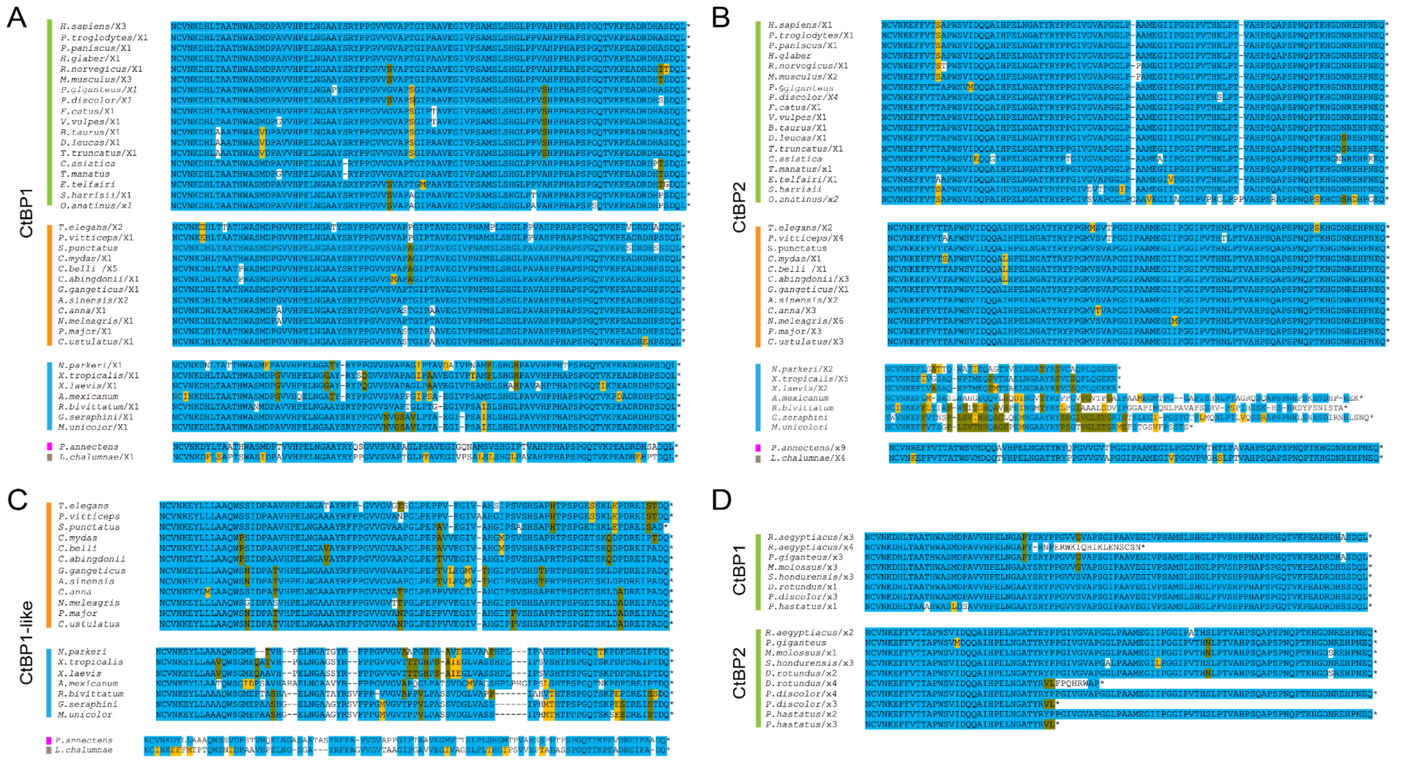


**Figure S8. Sarcopterygii have conserved CtBP1 and CtBP2 CTDs, with variations in amphibian CtBP2. A)** CtBP1 CTD alignment of all Sarcopterygii species analyzed. For all panels, vertical colored bars indicate the same clades as in Figure 10. **B)** CtBP2 CTD alignment of all Sarcopterygii species analyzed indicates high conservation of the CTD, aside from amphibia, where shorter and more divergent variants are observed. **C)** Alignment of CtBP1-like sequences, which are found in all Sarcopterygii other than mammals. **D)** Bats are the only mammals found to have short and long versions of CtBP1 and CtBP2 CTDs.

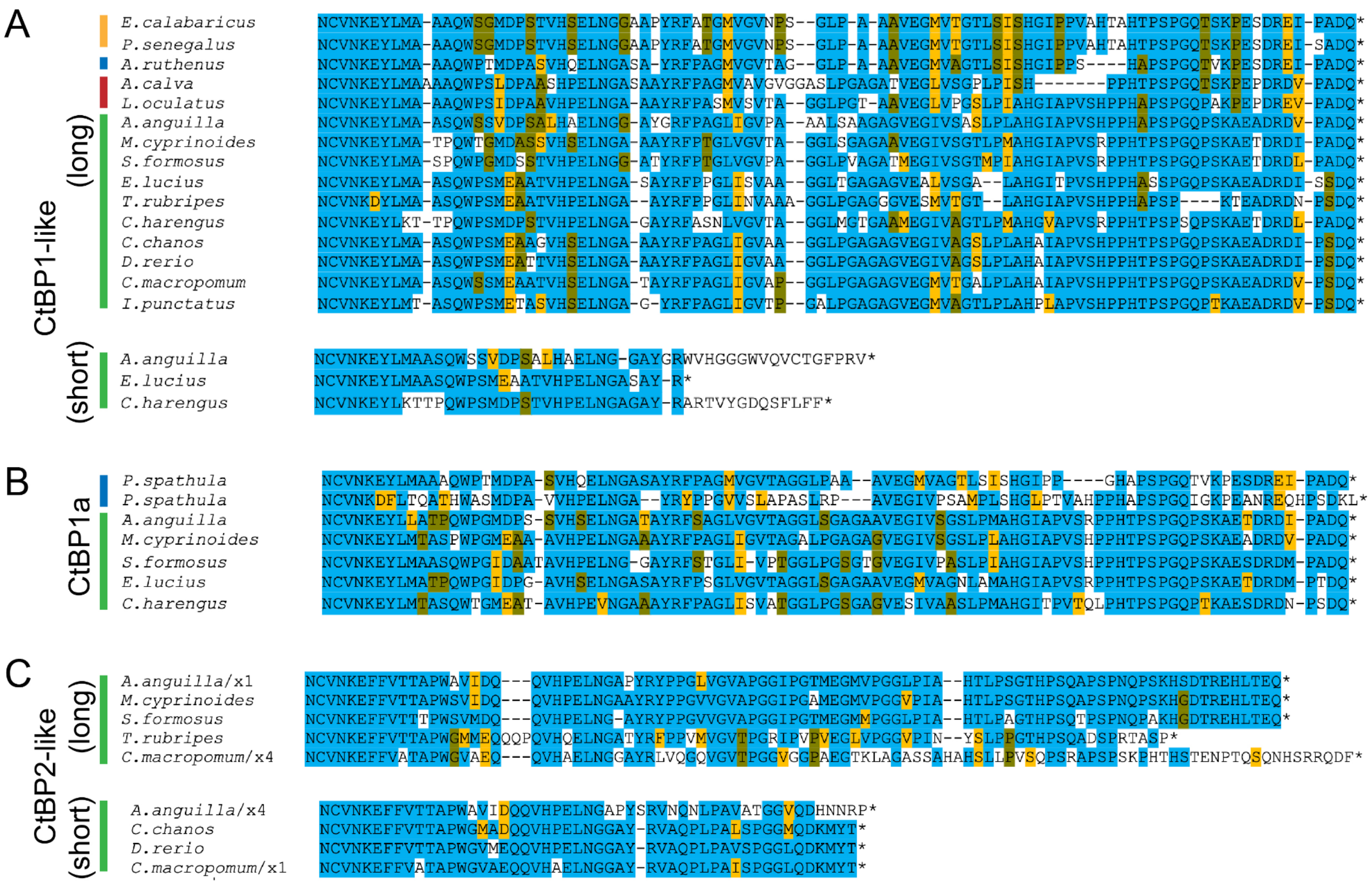

**Figure S9. Some Actinopterygii encode two additional CtBP paralogs. A)** Alignment of long isoforms of CtBP1-like. This protein is found in all Actinopterygii sampled. Short isoforms are present in certain Teleostei. **B)** Alignment of long isoforms of CtBP1a indicates this protein is conserved only in select Teleostei and Chondrostei. *P. spathula* seems to be the only species to have two versions of what we termed CtBP1a; it is unclear if one of them is indeed CtBP1-like. **C)** Alignment of CtBP2-like, which is found only in select Teleostei. Both long and short isoforms are encoded, and conserved across species. Vertical bars on the left correspond to the clades highlighted in Figure 11A. Due to mis-annotation of some of the genes in Actinopterygii, we renamed sequences from certain species (see Materials and Methods).

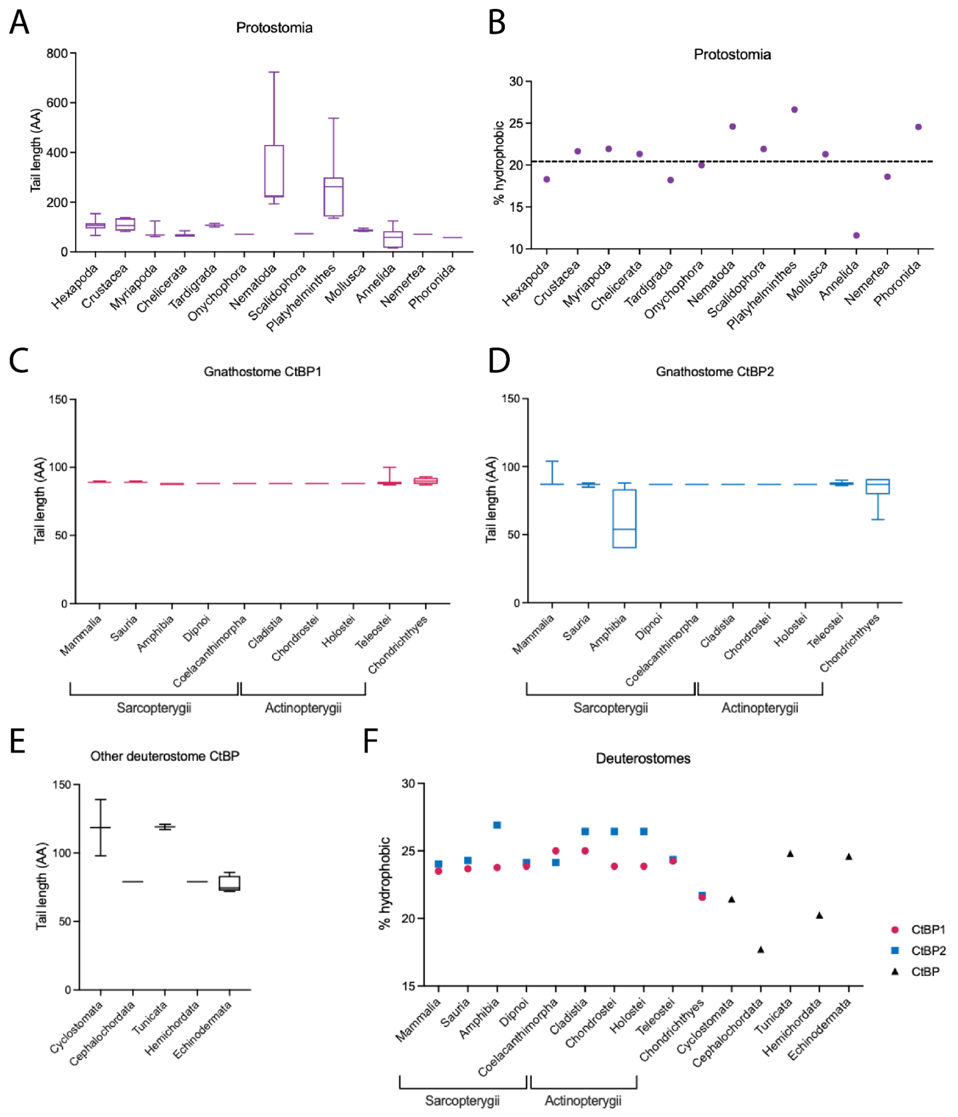


**Figure S10. Properties of CtBP CTDs in Bilateria. A)** CTD length (in Amino Acids, AA) across Protostomia. Average CTD lengths are similar except in divergent nematode and platyhelminth lineages, where some are over 500 AA. The leech family within annelids are the only phylum to have lost the long CTD. Box and whisker plots are used to indicate the middle 50% of values (box) and range (whiskers from lowest to highest value). Horizontal line indicates the median value. For all plots, tail properties were determined by averaging the longest CTD of all species within each clade. **B)** Hydrophobic content of CTD in protostomes (average ~21%, indicated by dashed line). **C)** Length of CtBP1 CTD in gnathostomes spanning Sarcopterygii, Actinopterygii, and Chondrichthyes. The length has remained mostly unchanged. **D)** Length of CtBP2 CTD in gnathostomes indicates that amphibians experienced some diversification in tail length. **E)** Length of CtBP CTD in non-Gnathostome deuterostomes also averages ~100 residues in length, but is more variable. **F)** Proportion of hydrophobic residues in vertebrate CtBP1 and CtBP2 CTDs and invertebrate deuterostome CtBP CTDs. Most classes have an average of about 24% hydrophobic residues, while some experience diversification, such as those within the non-vertebrate phyla. Percent hydrophobicity was determined by calculating the number of M, I, V, L, F, Y, and W residues in the longest tail from each species, and averaging the hydrophobic content across each clade. Clades include 1 to 18 species in total.

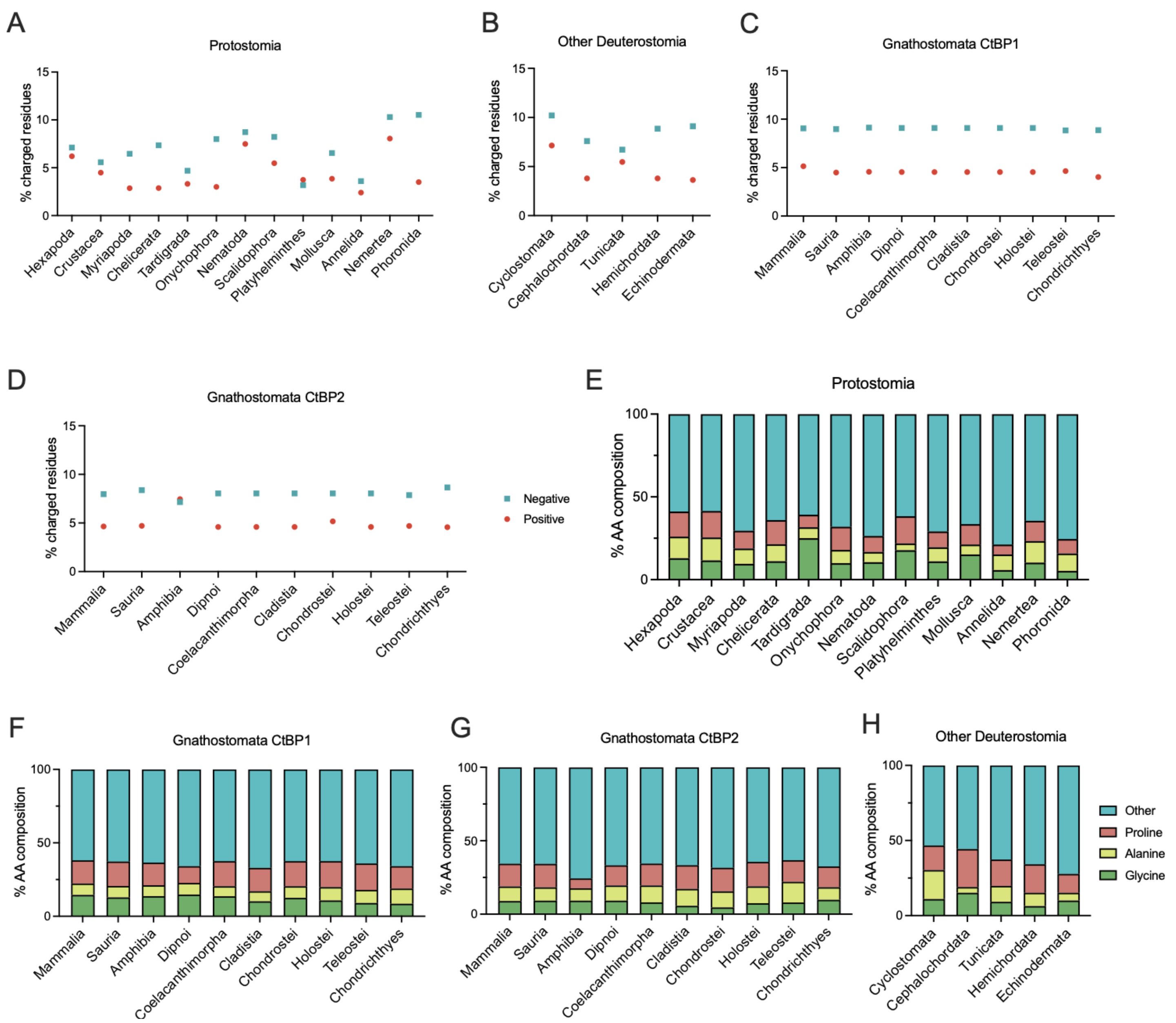


**Figure S11. Composition of amino acid residues in the CtBP CTD across Metazoa.** Composition of charged residues (positive: K, R; negative: D, E) in CtBP CTDs across **A)** Protostomia, **B)** non-gnathostome Deuterostomia, **C)** Gnathostomata CtBP1, and **D)** Gnathostomata CtBP2. Negatively charged residues are indicated by blue squares, and positively charged residues are indicated by red circles. We find that in deuterostomes, the charged residues are consistently making up under 15% of the CTD, while in protostomes there's more variability (6% in Annelida to 18% in Nemertea). AA composition of the CTD in **E)** Protostomia, **F)** Gnathostomata CtBP1, **G)** Gnathostomata CtBP2, and **H)** non-gnathostome Deuterostomia. Glycines are indicated in green, alanines in yellow, prolines in pink and all other residues in blue. We find that across Metazoa, most species have about 40% of their CTD made up of P, A, G residues, which are disorder-promoting. These analyses were performed by selecting one isoform of longest length from each species and averaging within each clade.

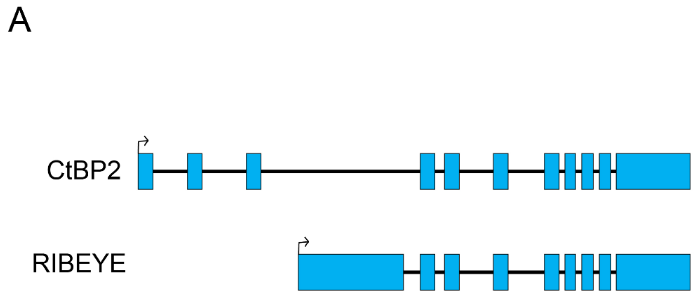


B

| | Species | NTD Length (AA) | % Conservation to human RIBEYE |
|---|---|---|---|
| Reference | *H.sapiens* | 572 | 100 |
| Mammalia | *M.musculus* | 575 | 79 |
| | *B.taurus* | 569 | 72 |
| | *P.discolor* | 556 | 65 |
| | *L.africana* | 577 | 72 |
| | *E.telfairi* | 553 | 65 |
| Sauria | *T.elegans* | 590 | 49 |
| | *P.vitticeps* | 598 | 53 |
| | *C.abingdonii* | 598 | 55 |
| | *A.sinensis* | 591 | 54 |
| | *P.major* | 573 | 53 |
| Amphibia | *X.tropicalis* | 607 | 45 |
| | *N.parkeri* | 605 | 45 |
| Coelacanth | *L.chalumnae* | 611 | 48 |
| Actinopterygii | *A.calva* | 621 | 45 |
| | *L.oculatus* | 614 | 46 |
| | *P.spathula* | 615 | 41 |
| | *E.calabaricus* | 619 | 45 |
| Chondrichthyes | *A.radiata* | 614 | 45 |
| | *C.carcharias* | 613 | 48 |
| | *C.punctatus* | 601 | 47 |

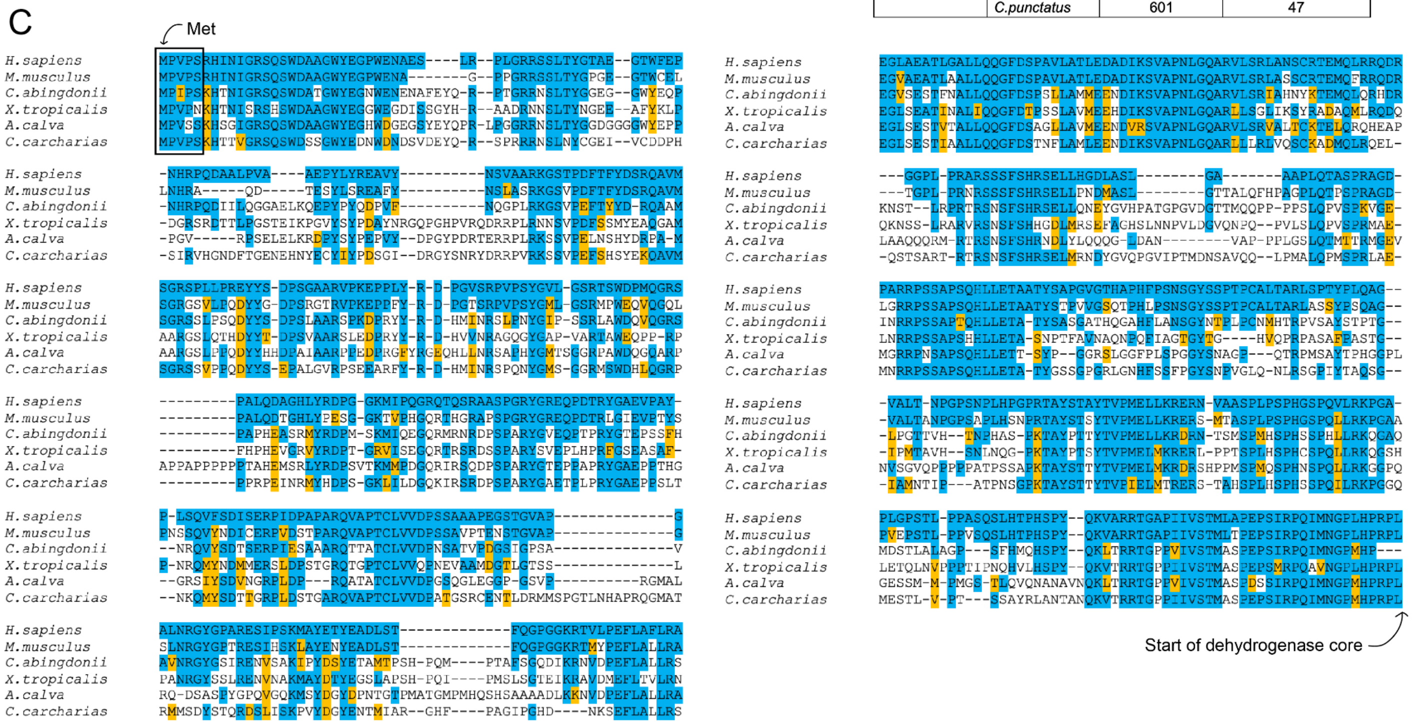


**Figure S12. N-terminal variations in CtBP2 are conserved. A)** Schematic of human CtBP2 and the RIBEYE variant with an extended NTD. Different transcriptional start sites are used to create a unique NTD. Blue boxes represent all exons (not to scale). **B)** Table indicating percent identity relative to the human sequence of the CtBP2 NTD in select gnathostomes. NTD lengths across Gnathostomata are consistent with the human RIBEYE variant length. Cyclostomata (not shown) do not have RIBEYE isoforms. **C)** Alignment of the NTD sequences from select gnathostomes. Blue highlight was used for the human RIBEYE sequence and for any residues that were completely conserved in the other species as compared to *H. sapiens*. Orange was used for chemical conservation compared to the human sequence. We find that the translational start site is highly conserved (boxed), and there are many blocks of conservation across the NTD. The sequence terminates with the RPL sequence, which is the start of the conserved dehydrogenase core.

A

| CtBP1 CTD modification | Effector | Impact | Reference |
|---|---|---|---|
| S422 (p) | HIPK2, JNK1 | CtBP degradation, apoptosis | Zhang *et al.* 2003; Zhang *et al.* 2005; Wang *et al.* 2006 |
| K428 (sm) | SUMO-1 | nuclear localization | Lin *et al.* 2003; Kagey *et al.* 2003 |

| CtBP2 CTD modification | Effector | Context | Reference |
|---|---|---|---|
| S365 (p) | Unknown | Found in ischemia | Mertins *et al.* 2014; Mertins *et al.* 2016 |
| T414 (p) | Unknown | Found in normal human liver tissue | Bian *et al.* 2013 |
| S428 (p) | HIPK2 | | Bian *et al.* 2013; Dewi *et al.* 2015 |

B

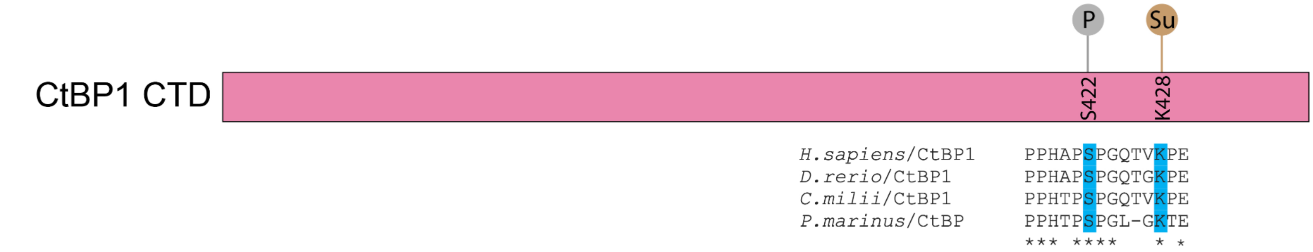


C

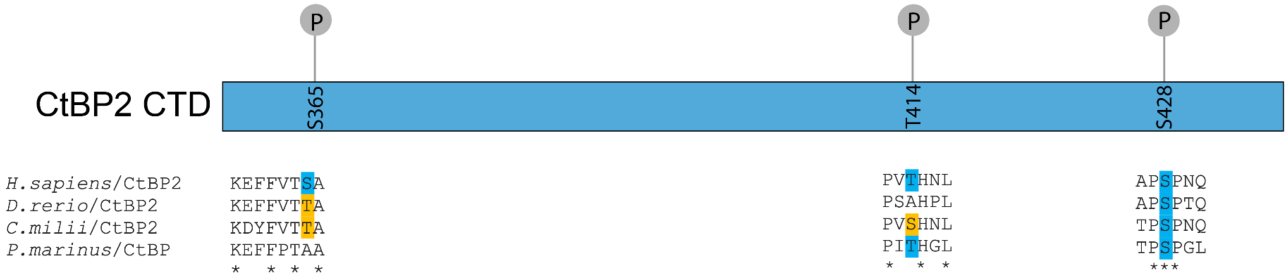


**Figure S13. Post-translational modifications of the human CtBP CTDs. A)** Chart summarizing experimentally validated PTMs on the *H. sapiens* CtBP1 and CtBP2 CTDs. **B)** Schematic of the *H. sapiens* CtBP1 CTD (89 residues). S422 and K428 are empirically determined phosphorylation and sumoylation targets, respectively. Both residues are conserved across vertebrates (blue highlight in alignment). **C)** Schematic of the *H. sapiens* CtBP2 CTD (87 residues). S365, T414 and S428 are empirically determined phosphorylation targets. Only S428 is conserved across vertebrates, while S365 and T414 show lower conservation.

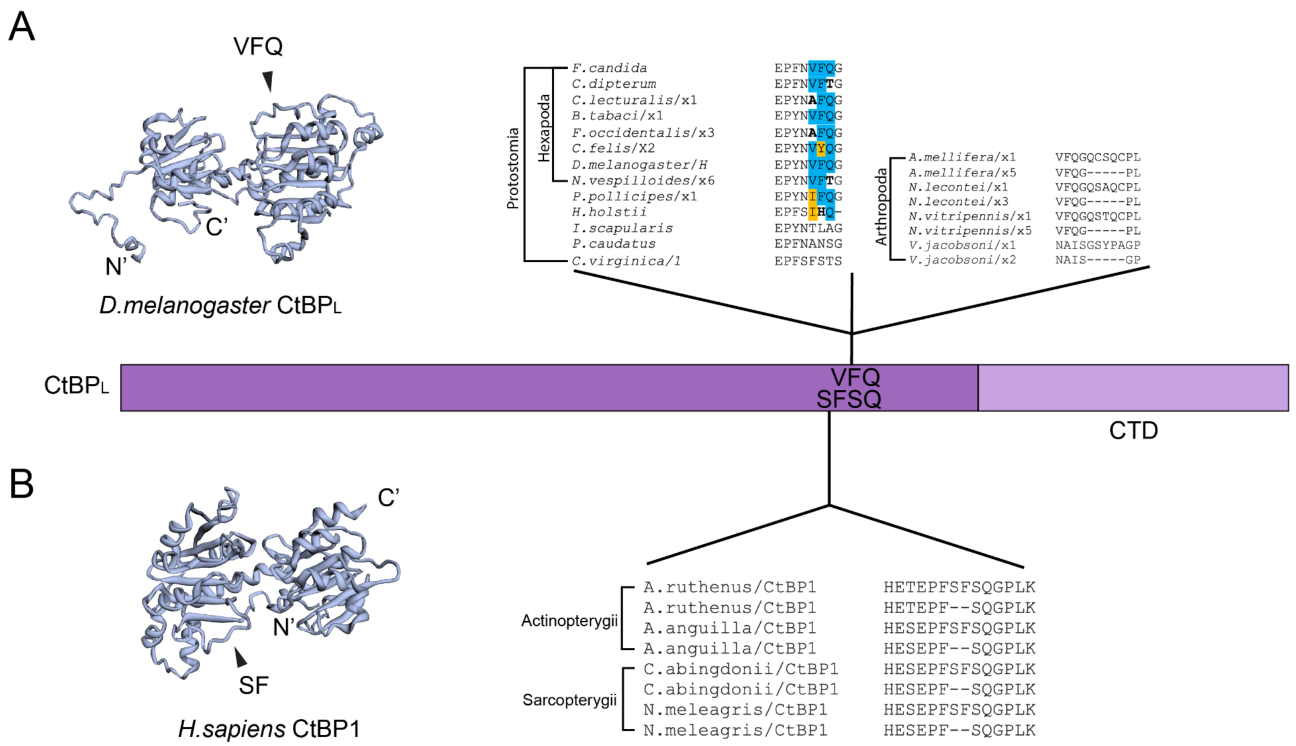


**Figure S14. Little variation in the CtBP dehydrogenase core. A)** Variation in the dehydrogenase core in Protostomia is limited to alternative splicing of the VFQ tripeptide in Diptera and alternative splicing of a 5mer in Arthropoda (alignments shown on right). This site is indicated on the Drosophila predicted structure on the left, and maps to an unstructured loop on the surface of the protein. Schematic of $CtBP_L$ is shown in purple, below. **B)** Variation in the dehydrogenase core in Deuterostomia is limited to alternative splicing of the SF motif in Actinopterygii and Sarcopterygii CtBP1 (alignment shown on right). This site is indicated on the human predicted structure on the left, and maps to an unstructured loop on the surface of the protein. These conserved alternative splicing events map to the same region of the core, near the CTD.
