## Supplemental File 2 for "The cynosure of CtBP: evolution of a bilaterian transcriptional corepressor"

**Bash script to calculate features of CtBP CTDs:**

```
# Remove the asterisk that is located at the end of some sequences
sed -i s/\"//g sequences_csv.csv

# Save the taxonomic classification field
awk -F ',' '{print $1}' sequences_csv.csv  | sed 's/ /_/g' > taxa.tmp

# Save the name of sequences
awk -F ',' '{print $2}' sequences_csv.csv  | sed 's/ /_/g' > names.tmp

# Save the length of the sequence for each field
awk -F ',' '{ print length($3) }' sequences_csv.csv > lengths.tmp

# Save the counts for the hydrophobic residues
awk -F',' '{print $3}' sequences_csv.csv | sed 's/[^(M,V,I,L,W,Y,F)]//g' | awk '{ print length }' >
hydrophobic.tmp

# Save the percentage of Hydrophobic residues for each sequence
paste -d"," hydrophobic.tmp lengths.tmp > 2c.tmp
awk -F',' '{print ($1/$2)*100}' 2c.tmp > p_hydrophobic.tmp

# Save the counts for the positive residues
awk -F',' '{print $3}' sequences_csv.csv | sed 's/[^(K,R)]//g' | awk '{ print length }' > positive.tmp

# Save the percentage for the positive residues
paste -d"," positive.tmp lengths.tmp > 2c.tmp
awk -F',' '{print ($1/$2)*100}' 2c.tmp > p_positive.tmp

# Save the counts for the negative residues
awk -F',' '{print $3}' sequences_csv.csv | sed 's/[^(D,E)]//g' | awk '{ print length }' > negative.tmp

#Save the percentage for negative residues
paste -d"," negative.tmp lengths.tmp > 2c.tmp
awk -F',' '{print ($1/$2)*100}' 2c.tmp > p_negative.tmp

# Save the counts for the number of glycines
awk -F',' '{print $3}' sequences_csv.csv | sed 's/[^(G)]//g' | awk '{ print length }' > glycines.tmp

# Save the percentage of glycines
paste -d"," glycines.tmp lengths.tmp > 2c.tmp
awk -F',' '{print ($1/$2)*100}' 2c.tmp > p_glycines.tmp

# Save the counts for the number of alanines
```

```
awk -F',' '{print $3}' sequences_csv.csv | sed 's/[^A]//g' | awk '{ print length }' > alanines.tmp

# Save the percentage of alanine
paste -d"," alanines.tmp lengths.tmp > 2c.tmp
awk -F',' '{print ($1/$2)*100}' 2c.tmp > p_alanines.tmp

# Save the counts for the number of prolines
awk -F',' '{print $3}' sequences_csv.csv | sed 's/[^P]//g' | awk '{ print length }' > prolines.tmp

# Save the percentage of proline
paste -d"," prolines.tmp lengths.tmp > 2c.tmp
awk -F',' '{print ($1/$2)*100}' 2c.tmp > p_prolines.tmp

# Paste all the values in a single one
paste -d ',' taxa.tmp names.tmp lengths.tmp hydrophobic.tmp p_hydrophobic.tmp positive.tmp
p_positive.tmp negative.tmp p_negative.tmp alanines.tmp p_alanines.tmp glycines.tmp
p_glycines.tmp prolines.tmp p_prolines.tmp  > table.csv

# Remove temporal files for a cleaner execution
rm *tmp
```
